# Interspecies synergism and antagonism induce differential and potentially exploitable susceptibility to various classes of antibiotics in a wound-like polymicrobial community

**DOI:** 10.64898/2026.04.28.721396

**Authors:** Caroline Laughlin-Black, Viviann Robles, Sabrina Wilson, Allie C. Smith, Catherine A. Wakeman

## Abstract

Chronic wounds are persistent and difficult to treat. Often this is because they are colonized by polymicrobial communities which contribute to changes in antimicrobial susceptibilities, making these infections harder to effectively clear. We explored the role a community can play in individual members’ survival when challenged by antibiotics, specifically looking at a community consisting of *Staphylococcus aureus*, *Pseudomonas aeruginosa*, *Enterococcus faecalis*, and *Acinetobacter baumannii*. Our data shows that communities can contribute to both increases and decreases in susceptibilities depending on the species and the antibiotic. The changes in susceptibilities can be due to interspecies cooperation or competition, with identifiable mechanisms. We also demonstrated that current antimicrobial susceptibility testing (AST) methods used in hospitals, which focus on determining the minimum inhibitory concentration (MIC) via determination of visible turbidity breakpoints, are not able to truly indicate the clearance of bacteria, as species can persist in higher antibiotic concentrations after visible turbidity is gone. To combat decreases in antibiotic susceptibilities contributed to by the community, we used our data from individual antibiotics to determine a potentially effective antibiotic combination, similar to combinatorial therapy used in hospitals to treat recalcitrant infections. Our data proved useful, as the combination of gentamicin and cephalexin was able to overcome polymicrobial synergism and clear the desired bacteria. This demonstrates that it is possible to determine effective antibiotic treatments for polymicrobial infections, whether they be combinatorial in nature or not. One simply must account for the role of the community in order to prescribe the most effective treatment.

## Introduction

Current antimicrobial susceptibility testing (AST) methods employed in hospitals focus on determining effective treatment for one causative agent of disease. If bacterial, antibiotic susceptibility is determined based on visible turbidity minimum inhibitory concentrations (MICs) (1, 2). These methods fail to account for both polymicrobial synergism as well as bacterial persisters surviving even after visible turbidity is lost. In order to better treat patients, one must establish the role the community plays in changes to individuals’ antimicrobial susceptibilities, as it has been shown that interactions in polymicrobial communities have effects on antibiotic susceptibilities (3). Our research focuses on a four species community consisting of *Staphylococcus aureus*, *Pseudomonas aeruginosa*, *Enterococcus faecalis*, and *Acinetobacter baumannii*. These pathogens are known to colonize chronic wounds, are ESKAPE pathogens, and are some of the leading causes of nosocomial infections (4, 5). They are also known to co-infect as communities, and a spectrum of interactions between these microorganisms ranging from competition to cooperation has already been characterized in various growth contexts.

Interactions between *S. aureus* and *P. aeruginosa* are well documented *in vitro*, in animal models of infection, and in human infections (6–9), with studies showing that coculture influences their individual antimicrobial susceptibilities (10, 11). Antibiotic resistant strains of *S. aureus* and *E. faecalis* have been shown to inhibit infection clearance in patients (12, 13), with species interactions leading to the formation of a more robust biofilm (14) while conversely, nonresistant strains of *E. faecalis* can display antagonism towards bystander *S. aureus* via the type VII secretion system (15). *S. aureus* and *A. baumannii* have been shown to antagonize each other in coculture (16) yet have been shown to colonize diabetic patients together (17) and work together to combat antibiotics like meropenem (18, 19). *P. aeruginosa* and *E. faecalis* are known to form dual-species biofilms together (20, 21) with *E. faecalis* contributing to *P. aeruginosa*’s antibiotic recalcitrance, especially to aminoglycosides (22, 23). *P. aeruginosa* is a known antagonist of *A. baumannii* (24), yet quorum sensing can allow for co-production of biofilms and increased populations of persister cells in response to oxidative stress in *A. baumannii* (25, 26). In many of the examples highlighted above, the specific culture conditions determined whether communities of bacteria would cooperate or compete (27–29).

Each of the above interactions focused on the interaction of just two species. However, infections can be even more complex and all of our species of interest can be found in a variety of combinations in sites of infection (30), with these different combinations having potentially different impacts on individual antimicrobial susceptibilities. Previous research conducted by our group showed that the antimicrobial susceptibility of these four organisms changes when they are grown in a community (31) (Supplemental Figures A1-A9). It also demonstrated that there are still populations of viable cells present even after visible turbidity is lost (Supplemental Figures A11-A14). Our recent research experiments focus on determining the mechanisms driving individual changes to antimicrobial susceptibilities when these four microorganisms are grown in a community as compared to monomicrobial suspension. The use of selective/differential media allows us to determine colony-forming unit (CFU) counts for each species, giving us a better depiction of the role the community plays in an individual’s survival. Based on our understanding of the mechanisms driving altered susceptibilities, we seek to exploit the relationships between community members in antibiotic prescription to make our antibiotic therapies more effective (32). Additionally, we strive to predict effective combinatorial therapies based on known species interactions driving susceptibilities and avoid combinations we know will result in treatment failure (33). Overall, by understanding the mechanisms driving altered susceptibilities within a community, it could be possible to exploit interactions between community members to achieve affective therapy use, improve patient infection clearing times, and promote effective antimicrobial stewardship.

## Materials and Methods

### Preparation of microbial cultures overnight

Approximately 5 colonies were obtained from a selective/differential plate using a sterile loop. Then the colonies were added to 5 mL of LB in a 125 mL glass flask. The culture was then incubated in a shaking incubator at 200 rpm and 37°C under ambient air conditions for 18 hours. After incubation, approximately 10^9^ CFU/mL were present in the overnight cultures.

### Preparation of bacterial inoculums

Per CLSI protocol, OD_625_ was measured using a spectrophotometer for each bacterial species (*S. aureus* ATCC 29213, *P. aeruginosa* ATCC 27853, *E. faecalis* ATCC 29212, and *A. baumannii* ATCC 19606) (1, 2). A 0.5 MacFarland standard equivalent concentration of OD_625_ 0.08 was created by diluting the bacteria in 1X PBS. To create the monomicrobial bacterial inoculums, 760 μL of 1X PBS was added to 40 μL of the MacFarland standard. In the polymicrobial inoculum, the 40 μL was divided by the number of species (10 μL per species since there were 4). Inoculums were also plated on selective/differential media in order to obtain inoculum CFU/mL.

### Competition assay setup

3 wells per species’ combination were filled with 90 μL of Cation-Adjusted Mueller Hinton Broth (CAMHB) in a 96-well plate. Species combinations consisted either of 2 or 3 species. The 40 μL inoculum was divided evenly among the 2-3 species (either 20 μL for each species when 2 were present in the community, or 13 μL per species when 3 were present in the community). 10 μL of the appropriate inoculum was added to each well. Competition assays were incubated for 18 hours before being diluted and plated in the same way as the AST panels, as described below.

### Antibiotic preparation and inoculation of antibiotic susceptibility testing (AST) panels

All wells in the first 3 rows of a 96-well plate were filled with 90 μL of CAMHB, as well as the first 6 wells in rows 4-6 (one row for growth control, one for vehicle control, and one for a negative contamination check). Using methods from the CLSI manual (1, 2), a 256 μg/mL storage stock was prepared for each desired antibiotic. 90 μL of the antibiotic stock was then added to the first well in rows 1-3. The wells were pipette-mixed and 90 μL was serially diluted across the rest of the wells in each row all the way to column 12 (the wells in column 1 contain 128 μg/mL of antibiotic with subsequent 1:2 dilutions leading to a final concentration of 0.06 μg/mL in the wells in column 12). 10 μL of the appropriate bacterial inoculum was added to each well in rows 1-3, as well as all the wells in the growth control and vehicle control rows.

### Diluting and plating the AST panels

After 18 hours, panels were pulled and read for visible turbidity. After filling a sterile pipetting reservoir with 1X PBS, 90 μL of 1X PBS was added to all wells in 96-well plates (as many as needed for each replicate in each column to have its own row of 8 when the panel is turned on its side). 10 μL of each well was added to the first well in a row (1:10 dilution), followed by serial dilutions of 10 μL into 90 μL all the way out to 10^-8^ (to save media, the monomicrobial panels only were diluted and plated at concentrations 3 wells above and 5 wells below the visible turbidity breakpoint). 5 μL of each diluted well was then plated onto selective/differential media in order to obtain colony counts (mannitol salt agar for *Staphylococcus aureus*, *Pseudomonas* isolation agar for *Pseudomonas aeruginosa*, bile esculin agar with azide for *Enterococcus faecalis*, and Leeds agar for *Acinetobacter baumannii*). After incubating 18-24 hours, the countable dilution factor was determined, and the CFU/mL for each well in the original AST panel was calculated.

### Data analysis for the AST panels and competition assays

Outliers were removed via outlier analysis in Microsoft Excel. Analysis for each set of individual antibiotic data and competition assays were performed via Mann Whitney test via GraphPad.

Significant points are denoted with an asterisk on the antibiotics’ graphs.

### Antibiotic preparation and dilution for the checkerboard assays

100 μL of CAMHB was added to each well in a 96-well plate. Using CLSI methods (1, 2), a storage stock with a concentration of 256 μg/mL was created for each antibiotic. 100 μL of the stock was added to all wells in the first column of the 96-well plate for antibiotic A. 100 μL was then diluted into the second column, and so on so that 1:2 serial dilutions were performed across the rest of the wells in the panel all the way to column 11 (column 1 had a concentration of 128 μg/mL and column 11 had a concentration of 0.125 μg/mL. The same process was repeated for antibiotic B in another 96-well plate.

### Checkerboard assay setup

Forty-five μL of each antibiotic was added at one concentration higher than desired to each well in 2 empty 96-well plates (one antibiotic on the x-axis and the other on the y-axis). Wells with a desired concentration of 128 μg/mL were filled with 45 μL of the 256 μg/mL stocks. Both the community and each species were given its own checkerboard. There were also 5 growth control wells (one for the community and one for each species), 5 vehicle control wells (phosphate buffer pH 6 for cephalexin), and 2 negative contamination check wells (one with and without vehicle). Except for negative contamination check wells, each well in the checkerboard was inoculated with 10 μL of the appropriate bacterial inoculum. After inoculation, all checkerboards were incubated at 37°C for 18 hours. Inoculums were diluted from 10^-1^ to 10^-8^ in 1X PBS and 5 μL was plated on the appropriate selective/differential media (inoculum values shown in Supplemental Table A.2).

### Diluting and plating the checkerboard assays

Checkerboards were pulled and read for visible turbidity after 18 hours of incubation. 2 dilutions were performed for the checkerboard assay. For the first dilution, 10 μL of checkerboard was added to 90 μL 1X PBS in each well of a 96-well plate (1:10 dilution). For the second dilution, 2 μL of the 1:10 dilution was added to 198 μL 1X PBS in a separate 96-well plate (1:100 dilution). 5 μL of the second dilution was plated on the appropriate selective/differential media and then incubated for 18-24 hours. Post incubation, monomicrobial and polymicrobial colony counts were obtained for each concentration of antibiotic in the checkerboards.

### Data analysis for the checkerboards

The CFU count for each replicate well, were averaged together. If the well was determined to be “to numerous to count” (TNTC), then 90 CFU was substituted as the upper limit of detection. To display the CFU counts, conditional formatting in Excel was used, so that higher CFU counts were indicated by darker coloration. The data in the graphs is representative of 3 replicates with “180” substituted for all “TNTC” wells when averaging replicates. Outliers were identified as wells separated from all other wells with growth by at least 1 column or row of zeroes and were removed from the graphs under the assumption that they represented cross contamination or genetic mutation allowing for obtainable counts. Statistical analysis for checkerboard assays was performed using an unpaired t-test with Welch correction. Antibiotic synergism or antagonism was determined using the equations outlined in a previous publication (33).

### Gentamicin AST with the addition of exogenous heme

Forty-five μL of gentamicin was added at one concentration higher than desired to each well in a 96-well plate (3 rows for AST). Wells with a desired concentration of 128 μg/mL were filled with 45 μL of the 256 μg/mL stock. 45 μL of CAMHB was added to wells 1-6 of rows 4-5 for growth control and contamination check. 10 μL of the appropriate bacterial inoculum was added to each well in rows 1-3, as well as all the wells in the growth control. 45 μL of CAMHB with 40 uM heme was then added to each well for a final concentration of 20 uM heme per well. Panels were then incubated at 37°C for 18 hours, and then diluted and plated on selective and differential media as described above.

### Anaerobic E. faecalis AST

AST panels were set up as described above, and then incubated at 37°C for 18 hours in an anerobic chamber (oxygen PPM >0.01).

### Determining the mechanism of E. faecalis’ decreased susceptibility to cephalexin

First the most influential community member on *E. faecalis*’ survival was determined by using an AST panel in combination with selective/differential media as described above, but with the paired inoculums as described in the competition assay protocol. After determining the most influential community member, the likely mechanism was determined as follows. To test heat-labile compounds, community member cells were heat shocked in 98°C heat for 5 minutes before being added to the AST panel. To test the supernatant, community member cells were spun down via centrifuge for 5 minutes at 2000 rpm. The supernatant was then filtered using a 0.2 µm filter to remove any remaining cells before being added to the AST panel. For supernatants after exposure to antibiotics the organisms were grown in the presence of the antibiotic for 18 hours, and then spun and filtered before being added to the AST panel. To test live cells from the community member that could not directly interact with *E. faecalis*, a 96-well transwell plate was used with membrane filter size of 0.4 μm. All panels were diluted and plated as described above after 18-hour incubation at 37°C.

### Confirming the presence of a beta-lactamase using nitrocefin discs

2 5 mL cultures of *A. baumannii*, one containing 128 ug/mL cephalexin, were grown in Lysogeny Broth for 18 hours at 37°C and 150 rpm. After incubation, the supernatants were filtered and 10 uL were added to Thermo Scientific™ Nitrocefin Disks (3 discs per supernatant per replicate), and pictures of the discs were taken at 0 seconds, 30 seconds, 1 minute, 2 minutes, 5 minutes, 10 minutes and 30 minutes. ImageJ software was used to quantity the darker color of the discs over time.

## Results

Polymicrobial interactions are important factors in determining whether an individual will survive within its community. This is demonstrated with *P. aeruginosa*’s antagonism of *A. baumannii* leading to *A. baumannii*’s death whenever *P. aeruginosa* is present within the community (Figure 2.1). These interactions become even more important in the presence of environmental stressors such as antibiotics, as community dynamics can influence antimicrobial susceptibilities (Table 2.1) (Supplemental Figures A.1-A.9). Cooperation/synergism between species can lead to recalcitrance against antibiotics, while competition/antagonism between species can lead to increased susceptibility (3, 7–34). It is important that we acknowledge that susceptibility changes can occur in both directions, with organisms becoming both more or less susceptible, in order to fully understand the role of communities in effective treatment of infections.

**Table 1.** Overall antibiotic trends show that polymicrobial communities can play a role in increasing or decreasing antibiotic susceptibilities.^a^.

| **Antibiotic Name** | **Changes to Antimicrobial Susceptibility (Aerobic)** |
| --- | --- |
| Gentamicin | ↑ *E. faecalis* |
| Tetracycline | ↑ *S. aureus* at lower concentrations<br>↑ *A. baumannii*<br>↓ *S. aureus* at higher concentrations (possible persisters) |
| Rifampin | ↑ *S. aureus*<br>↑ *A. baumannii* |
| Ceftazidime | ↓ *P. aeruginosa* |
| Cephalexin | ↑ *S. aureus*<br>↑ *A. baumannii*<br>↓ *E. faecalis* |
| Ciprofloxacin | ↑ *S. aureus*<br>↓ *P. aeruginosa* |
| Polymyxin B | ↑ *E. faecalis* |
| Vancomycin | ↓ *E. faecalis*<br>↓ *A. baumannii* |
| Augmentin<br>(Amoxicillin + Clavulanate) | ↑ *S. aureus*<br>↓ *E. faecalis* |

<sup>a</sup>Blue arrows pointing up represent an increase in susceptibility to the antibiotic by that specific species while red arrows pointing down represent a decrease in susceptibility to the antibiotic by that specific species. As seen above, susceptibilities can change for each species because of the community and can differ based on the antibiotic (the same species can see an increase in susceptibility to one antibiotic and a decrease in susceptibility to another all because of the same community).

**Figure 1.**
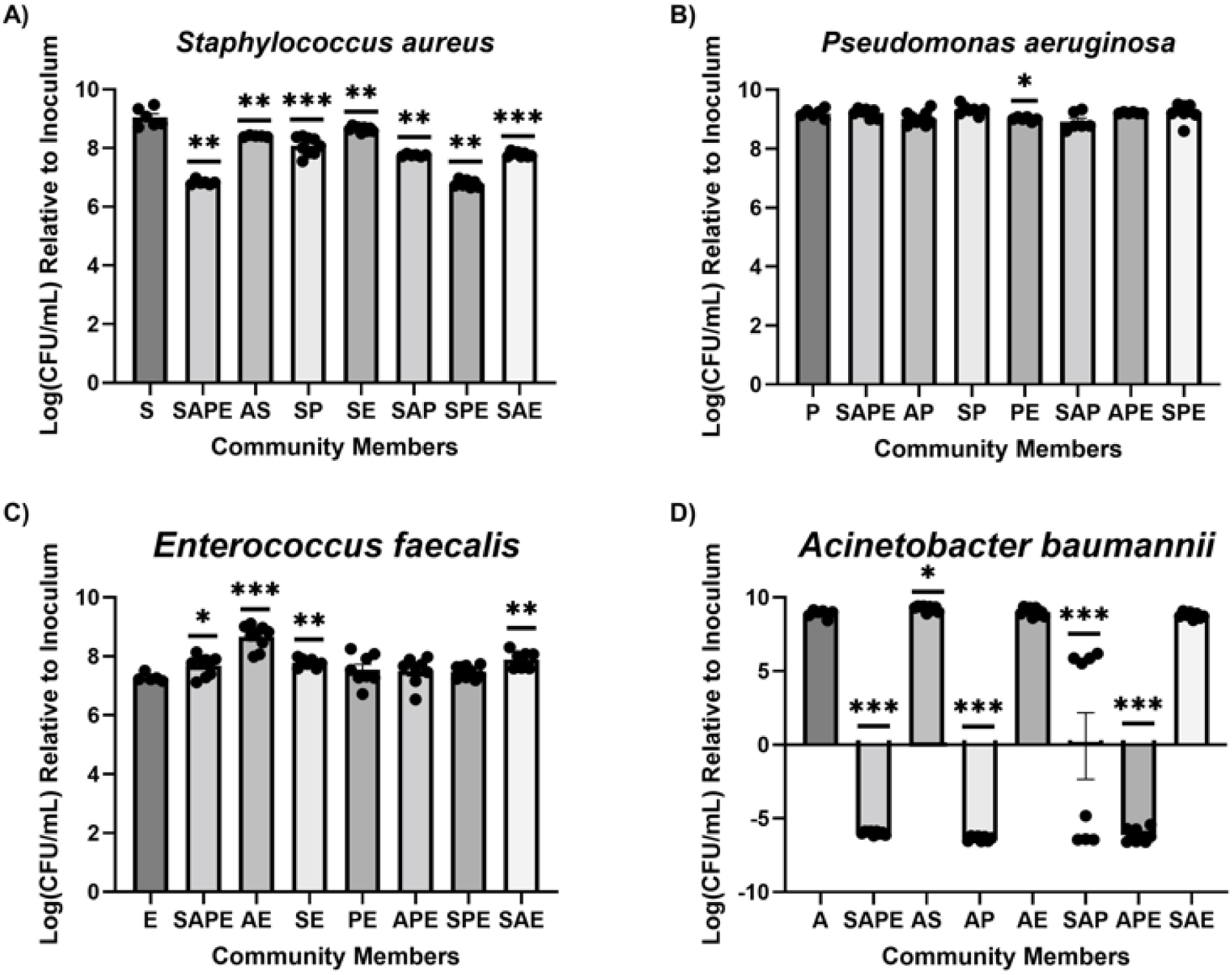
Polymicrobial Competition Assays show changes in growth levels dependent on interspecies competition or synergism. Inoculum CFUs are recorded relative to the inoculum. “S” represents *S.* aureus, “P” represents *P. aeruginosa*, “E” represents *E. faecalis*, and “A” represents *A. baumannii*. Combination of letters, such as “SAPE” represent communities and list the first letter of each member (so “SAPE” would contain all 4 members of the polymicrobial community, while “AS” would contain just *A. baumannii* and *S. aureus*). Phenomenon worth noting is *S. aureus*’ decrease in growth whenever present in a community with *P. aeruginosa*, or with both *E. faecalis* and *A. baumannii*. *P. aeruginosa* does not seem to prefer certain communities over others, simply growing at the same rate each time. *E. faecalis* appears to do better when it is grown in community rather than by itself. *A. baumannii* does poorly in any community containing *P. aeruginosa* due to antagonism (24). Asterisks represent significance as determined by a Mann Whitney test.

**Figure 2.**
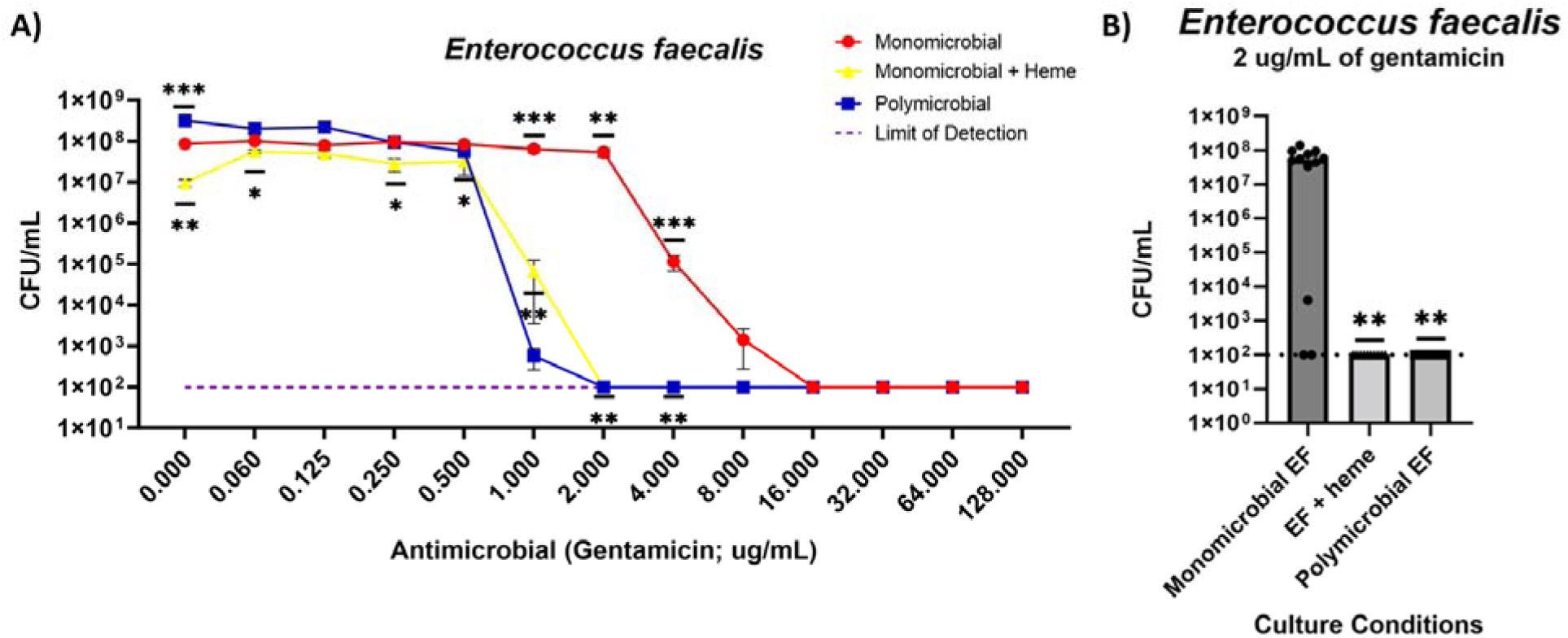
*Enterococcus faecalis* shows increased susceptibility to gentamicin when grown in the polymicrobial condition due to heme cross-feeding. *E. faecalis* displayed increased susceptibility to gentamicin in the polymicrobial community (1 μg/mL to 8 μg/mL) due to heme cross-feeding. Addition of exogenous heme to monomicrobial *E. faecalis* induced the same susceptibility trend (shown in yellow). All lines are indicative of data averaged from 9-12 technical replicates across 3-4 biological replicates (3 technical replicates per biological). Asterisks represent significance as determined by a Mann Whitney test.

Sharing of community resources also influences susceptibility to antimicrobials. *E. faecalis*, especially, cannot make heme on its own, however, it can acquire it from other community members or from the host environment. Previous studies have demonstrated that heme acquisition by *E. faecalis* can enhance the organism’s fitness in infection (35, 36), and help it survive against antimicrobials, (such as hydrogen peroxide via mechanisms such as activation of heme-dependent catalases (37)). Our research suggests that heme acquisition by *E. faecalis* from its fellow community members sensitizes the bacterium to the antibiotic gentamicin. This sensitization phenotype from the polymicrobial community was replicable with monomicrobial *E. faecalis* with the addition of exogenous heme (Figure 2.2). We believe this sensitization is due to heme cross-feeding leading to alteration of the bacterium’s proton motive force. The antibiotic gentamicin requires aerobic respiration to enter bacterial cells where it can reach its target – the 30S ribosome. *E. faecalis* typically performs fermentation, but the availability of heme can induce aerobic respiration (14). This induction of respiration increases *E. faecalis*’ susceptibility to gentamicin as its proton motive force has changed, allowing the oxygen-dependent transport of gentamicin into the cell where it can reach the ribosome, thus sensitizing the bacterium to the antibiotic. However, this phenomenon only occurs in aerobic environments, where respiration is favorable. When *E. faecalis* was grown within the community in anaerobic conditions, this sensitization no longer occurred (Supplemental Figure A.10), further supporting that the community-driven change in antibiotic susceptibility is due to activation of aerobic respiration by heme crossfeeding. These results not only demonstrate that shared resources within a community can alter antimicrobial susceptibilities, but also that changes in susceptibility are not always dependent solely on the presence of other community members, but rather also the environment the community is in. The infection environment (not just other community members, but also the environmental setting itself) is crucial to take into account when determining AST for effective treatment prescription (38).

While *E. faecalis*’ sensitization to the antibiotic because of the presence of community members was seen with gentamicin, we also saw decreased susceptibility in the same organism to the antibiotic cephalexin when grown in the presence of the community. We sought to determine which community member(s) were needed to induce this decreased susceptibility phenomenon. After experimenting with all combinations of community members, it was determined that *A. baumannii* contributed to the greatest *E. faecalis* survival (with *P. aeruginosa* also contributing, but at a lesser level) (Figure 3). In order to begin to determine the mechanism of recalcitrance, we used a transwell plate to determine whether *A. baumannii* and *E. faecalis* needed to physically interact to induce the recalcitrance observed. The transwell plate demonstrated similar survival levels of *E. faecalis* as compared to the results observed with *A. baumannii* in a normal 96-well plate (no transwell membrane) (Figure 2.3). Therefore, it was reasoned that the mechanism must not be dependent on physical interactions between the 2 organisms, but rather something released from *A. baumannii* that is small enough to cross the membrane. However, exposure to *A. baumannii*’s supernatant without cephalexin present did not induce the recalcitrant phenomenon, therefore, it was determined the mechanism is likely dependent on the presence of the antibiotic. Therefore, we exposed *A. baumannii* to the highest concentration of cephalexin tested (128 µg/mL), filtered the cells from the supernatant, and added that to monomicrobial *E. faecalis* challenged with the same concentration of antibiotic.

**Figure 3.**
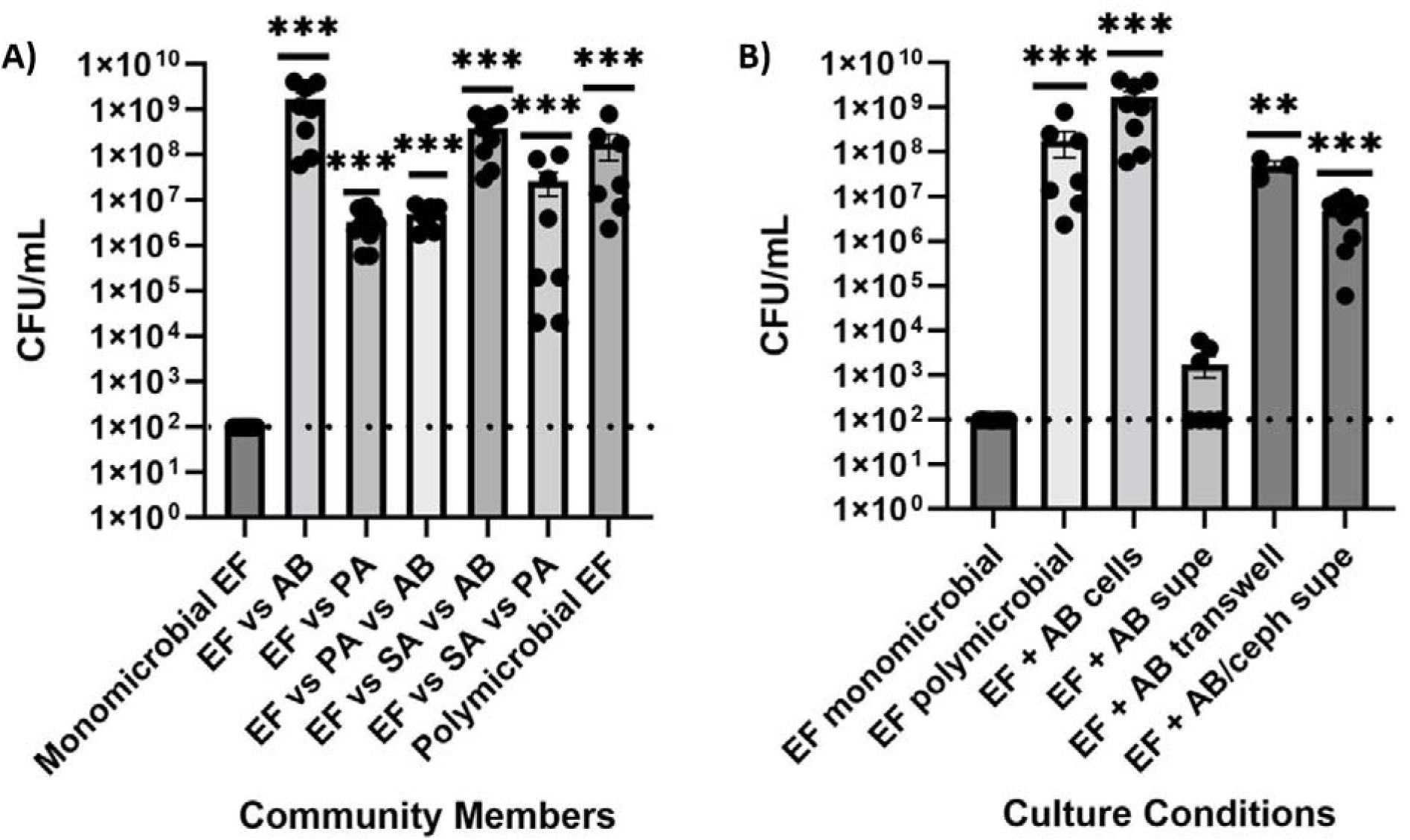
*Enterococcus faecalis* decreased susceptibility to cephalexin is likely due to a beta-lactamase produced by gram-negatives, especially *A. baumannii*. A. *E. faecalis* displayed decreased susceptibility to 128 µg/mL cephalexin in the presence of *A. baumannii* and *P. aeruginosa*. **B.** This is likely due to production of a beta-lactamase, as heat killing the cells removed the phenomenon, and using the filtered supernatant (supe) of *A. baumannii* after exposing the bacterium to the highest concentration of antibiotic (128 μg/mL) induced the decreased susceptibility phenomenon seen with live *A. baumannii* cells. Asterisks represent significance as determined by a Mann Whitney test.

**Figure 4.**
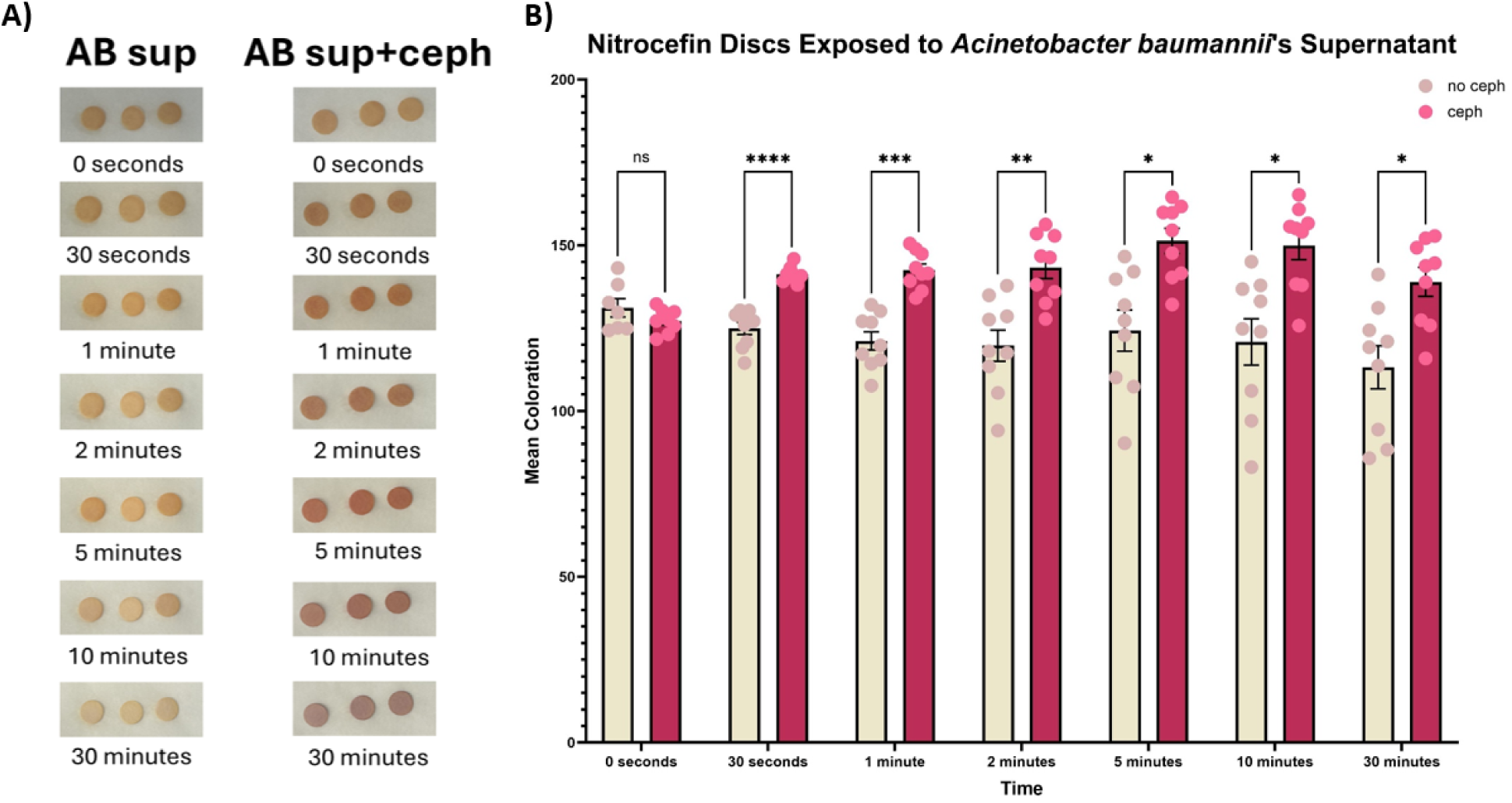
Beta-lactamase production by *A. baumannii* protects *E. faecalis* from cephalexin. **A.** A time course assay using nitrocefin discs demonstrated that discs inoculated with *A. baumannii*’s supernatant after cephalexin exposure turned red over time, while discs inoculated with supernatant with no cephalexin exposure stayed yellowish tan. **B.** Discs from both supernatants showed no significant difference in coloration at 0 seconds, but discs inoculated with supernatant post cephalexin exposure showed a significant increase in coloration at all other time points as compared to discs inoculated with supernatant with no cephalexin exposure. Asterisks represent significance as determined by a Tukey’s multiple comparisons test.

**Figure 5.**
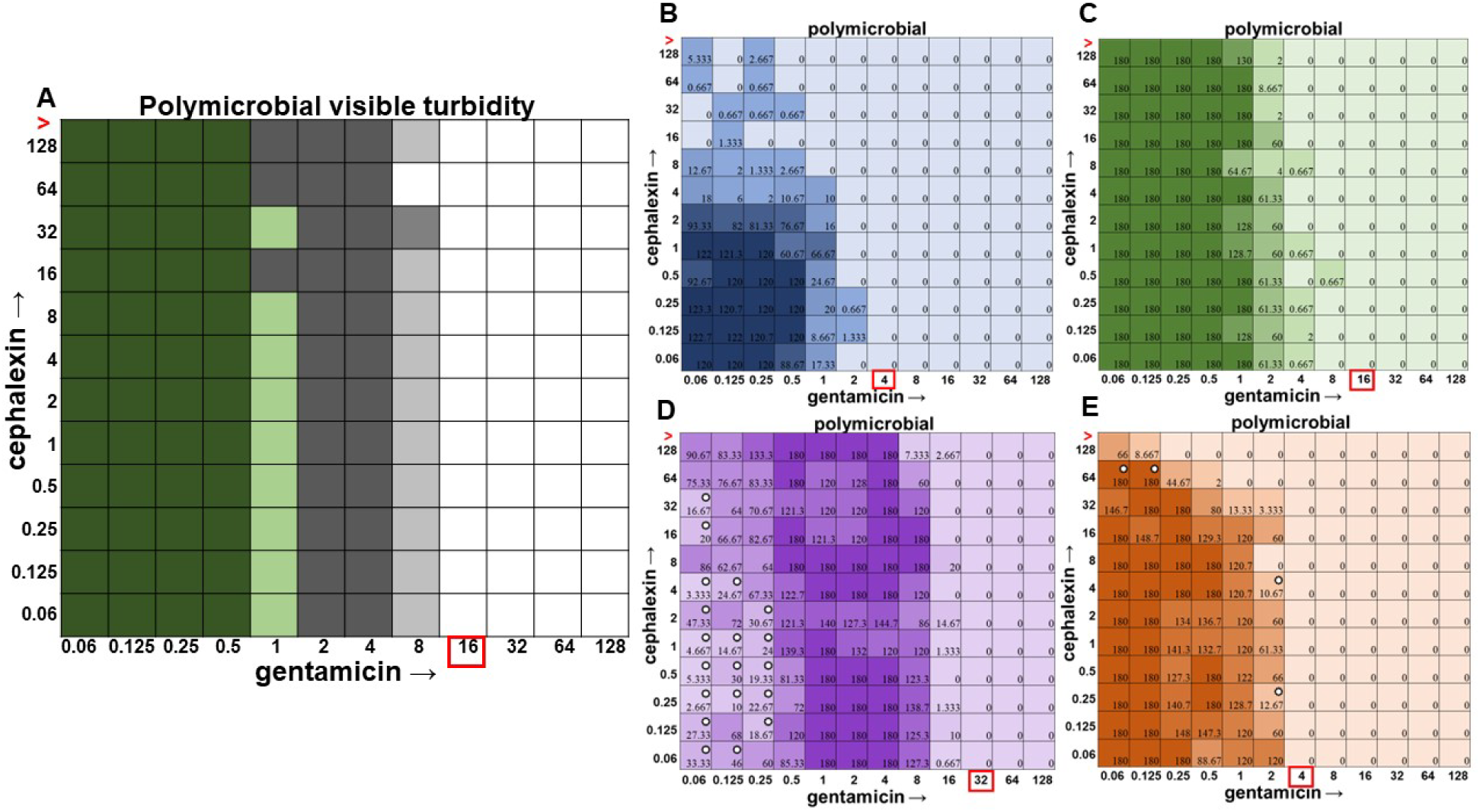
Checkerboard assay with the four species polymicrobial community. **A.** Visible turbidity of the polymicrobial community consisting of all 4 species. Data represents triplicate checkerboard assays with cephalexin and gentamicin. Green represents where *P. aeruginosa* pigmentation is present, while gray represents turbidity without green pigmentation. Dark coloration indicates that turbidity was visible in the well in all replicates, medium coloration indicates that turbidity was visible in 2 replicates, light coloration indicates that turbidity was visible in 1 replicate. **B-E.** CFU counts from each individual species when grown in the polymicrobial condition are shown on the right (**B.** *S. aureus* in blue, **C.** *P. aeruginosa* in green, **D.** *A. baumannii* in purple, and **E.** *E. faecalis* in orange). As shown above, bacteria are still present even after there is no longer any visible turbidity. Loss of *P. aeruginosa*’s green pigmentation is also associated with loss of interspecies competition, as shown by the increase in *A. baumannii*’s population as the pigment disappears from view. Data represent averages of triplicates performed on different days. Stars represent significance as determined by an unpaired t-test with Welch correction.

**Figure 6.**
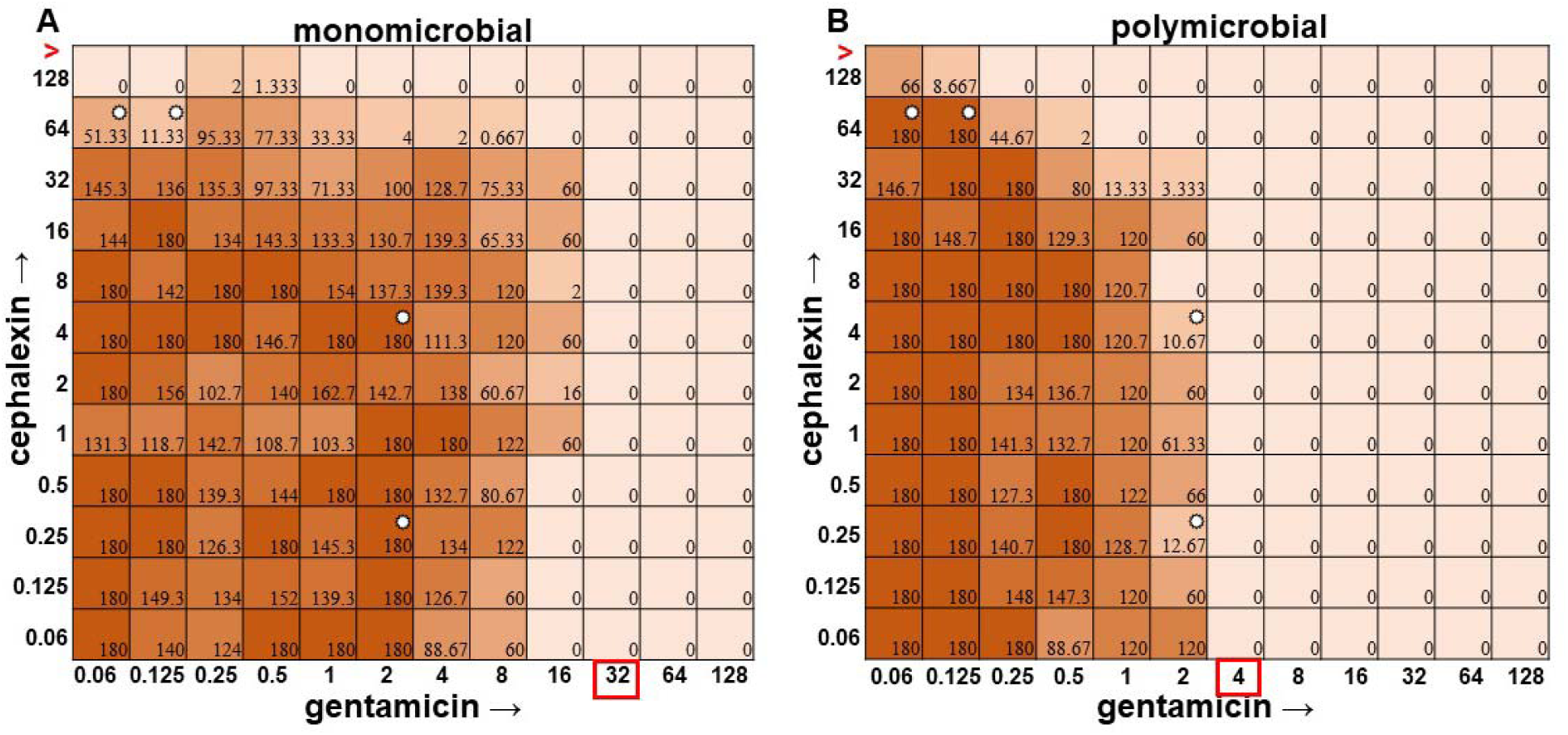
Monomicrobial versus polymicrobial checkerboards reveal that *E. faecalis* loses sensitivity to cephalexin in polymicrobial culture but gentamicin has an additive effect. CFU counts for *E. faecalis* ATCC 29212 when grown in the **A.** monomicrobial condition versus the **B.** polymicrobial condition. As shown above, *E. faecalis* becomes sensitized to gentamicin at 4 μg/mL, overcoming the lack of sensitivity to cephalexin at 128 μg/mL. Stars represent significance as determined by an unpaired t-test with Welch correction.

**Figure 7.**
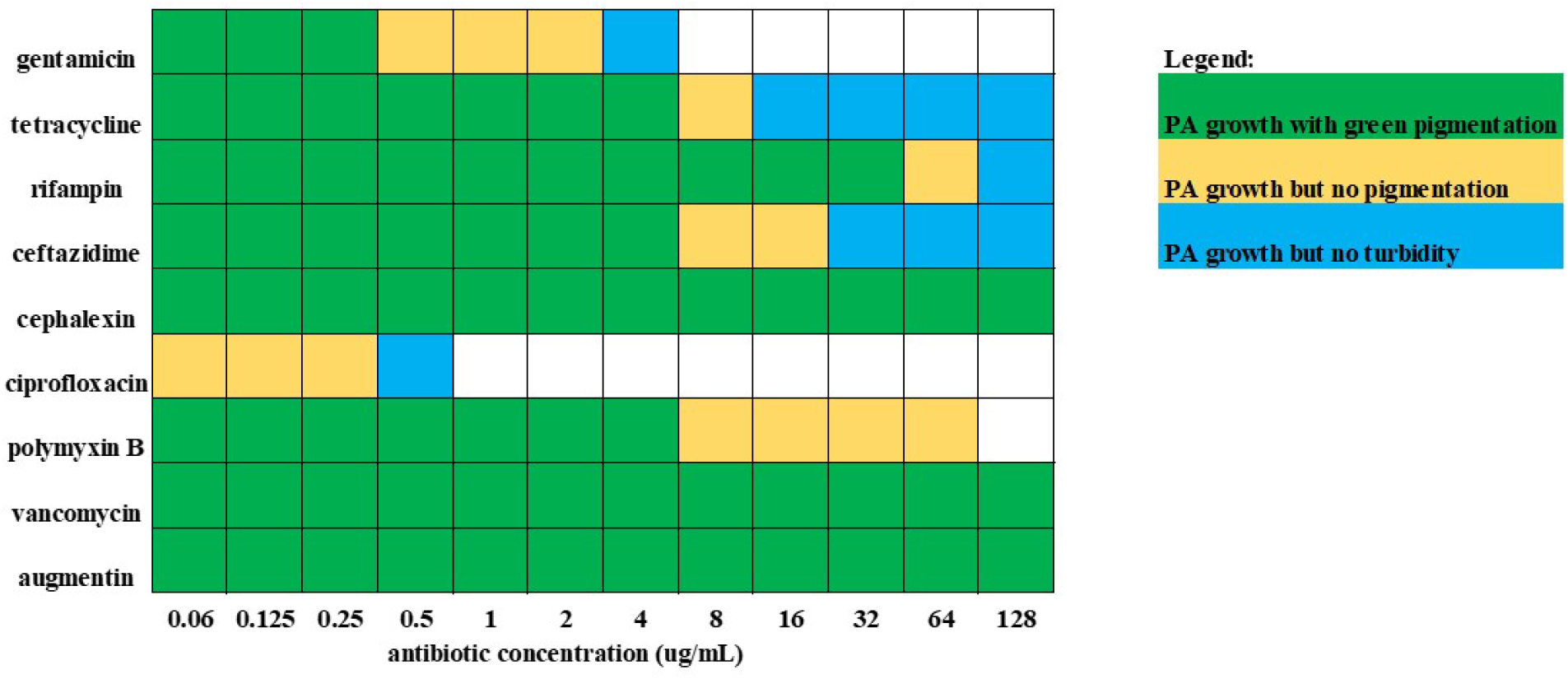
*Pseudomonas aeruginosa* visible turbidity in the polymicrobial condition is not indicative of the present of CFUs. Presence of pyocyanin pigment is not always seen with *P. aeruginosa* growth as it is population-dependent. Overall visible turbidity does not accurately depict where detectable levels of CFUs can be found. Often *P. aeruginosa* still has detectable levels of CFUs, even after visible turbidity is no longer seen by the naked eye.

This supernatant collected post antibiotic exposure recreated the recalcitrance observed previously, and so it was determined that the supernatant likely contains a compound capable of breaking down the antibiotic before it reaches *E. faecalis*, and only produced in high quantities in the presence of the antibiotic, such as a beta-lactamase (Figure 2.3). To confirm whether a beta-lactamase was present, nitrocefin discs were used and a time-course assay was performed to determine the level of color change over time for discs exposed to *A. baumannii*’s supernatant, with and without cephalexin exposure. The results demonstrate that a beta-lactamase is indeed present in *A. baumannii*’s supernatant post-cephalexin exposure, but not in the supernatant with no cephalexin exposure (or at least not at the level to be detected by the discs) (Figure 2.4). Therefore, the beta-lactamase produced by *A. baumannii* when cephalexin in present protects susceptible *E. faecalis* from death.

Based on the results obtained for gentamicin and cephalexin, we wanted to determine if the sensitization effect seen with gentamicin for *E. faecalis* within the polymicrobial community was enough to overcome the recalcitrance seen with cephalexin. We performed a checkerboard assay and calculated the antibiotic interaction using a previously described methodology (33).

We saw similar trends with the polymicrobial checkerboard as we saw with our single antibiotic data. *P. aeruginosa* continued to outcompete *A. baumannii*, which correlated nicely with visible pyocyanin production, which made sense as pyocyanin is known to help with bacterial competition (39) (Figure 2.5). We determined that the sensitization of *E. faecalis* to gentamicin from 32 μg/mL to 4 μg/mL was enough to overcome the recalcitrance to cephalexin (Figure 2.6). Therefore, the combination of gentamicin with cephalexin has an additive effect when treating *E. faecalis* within the polymicrobial community.

Another interesting phenomenon to note is the number of CFUs still present both in the checkerboard and in single antibiotic panels, even when no visible turbidity can be seen. Currently, hospitals often rely on visible turbidity readings to determine the MIC of antibiotics before prescribing them. However, as shown in Figures 2.5 and 2.7 (and Supplemental Figures A.11-A.14), species in a community often have detectable levels of CFUs at concentrations of antibiotic much higher than the breakpoint where loss of visible turbidity is seen. Just because there are no longer any bacteria visible to the naked eye or detected via spectrophotometer reads, does not mean that there are no bacteria present. These bacteria have been shown to persist even at higher concentrations of antibiotic, which could prove problematic when attempting to effectively treat patient infections.

## Discussion

The data from this study highlights the importance of understanding the role of bacterial community interactions for determination of bacterial infection dynamics. Polymicrobial communities can lead to either increases or decreases in susceptibilities to clinically relevant antibiotics. However, we have shown that if community-mediated decreases in antibiotic susceptibility are a concern, combinatorial therapies can be used to overcome polymicrobial synergism. While the polymicrobial community decreases *E. faecalis*’ susceptibility to cephalexin, an antibiotic used to treat gram-positive infections, the additive effect of gentamicin was enough to clear *E. faecalis* bacteria, even at lower concentrations of gentamicin. This demonstrates that while increases in antibiotic resistance (driven by genetics) and tolerance (driven by phenotypic adaptation) are a major concern in healthcare, the effective life of an antibiotic can be extended using planned combinatorial therapies that take into account the role of the community in infection.

One of the biggest shortcomings of current AST methods in hospitals is the use of visible turbidity to determine the MIC of an antimicrobial. As shown in our data, bacteria can persist at higher levels even after the loss of visible turbidity. These persisters put patients at risk for prolonged infection as the antimicrobial prescribed may not be enough to clear the infection completely (or may not be prescribed at the dosage needed to effectively treat the patient). While it may not be necessary to clear 100 percent of the bacteria in order to clear the infection, as the immune system can often take care of the bacteria once they drop below a certain level, leaving high concentrations of bacteria present, (such as *S. aureus* with cephalexin having levels of bacteria up to 10^7^ CFU/mL in the monomicrobial condition even after loss of visible turbidity (Supplemental Figure A.5),) could be dangerous to the patient could be dangerous to the patient, especially if they are immunocompromised. While visible turbidity readings are relatively quick to obtain and can tell us a lot about how bacteria are reacting to the antibiotic, they do not always provide the most accurate determination of MIC. It is imperative that we take that into account when attempting to prescribe antibiotics based on MICs obtained from visible turbidity readings.

Another shortcoming of current AST methods is the use of monomicrobial cultures determined to be the causative agent of disease. As demonstrated, the community can play a role in changing antimicrobial susceptibilities, for better or worse. Failing to take into account the role of the community can be dangerous, as polymicrobial communities are often associated with increased patient morbidity and mortality (3). Our research has demonstrated that the mechanisms of polymicrobial interactions can be determined and exploited in effective treatments. In order to effectively treat patient infections, the community the pathogen is in must be recognized as we attempt to prescribe antimicrobials that will clear the infection.

Combinatorial therapies can help by understanding the community’s role and designing therapies based on them can help extend the life of our current antibiotics, providing an opportunity to fight recalcitrant infections with tools we already possess.

## ACKNOWLEDGEMENTS

This research was funded by NIH/NIAID grant number R01 AI173686, and a TTU mid-career grant.

