## Supplemental Figures for "Interspecies synergism and antagonism induce differential and potentially exploitable susceptibility to various classes of antibiotics in a wound-like polymicrobial community"

**Supplemental Table A1. Average inoculum colony-forming units per milliliter and standard deviations for each community in the competition assays.**

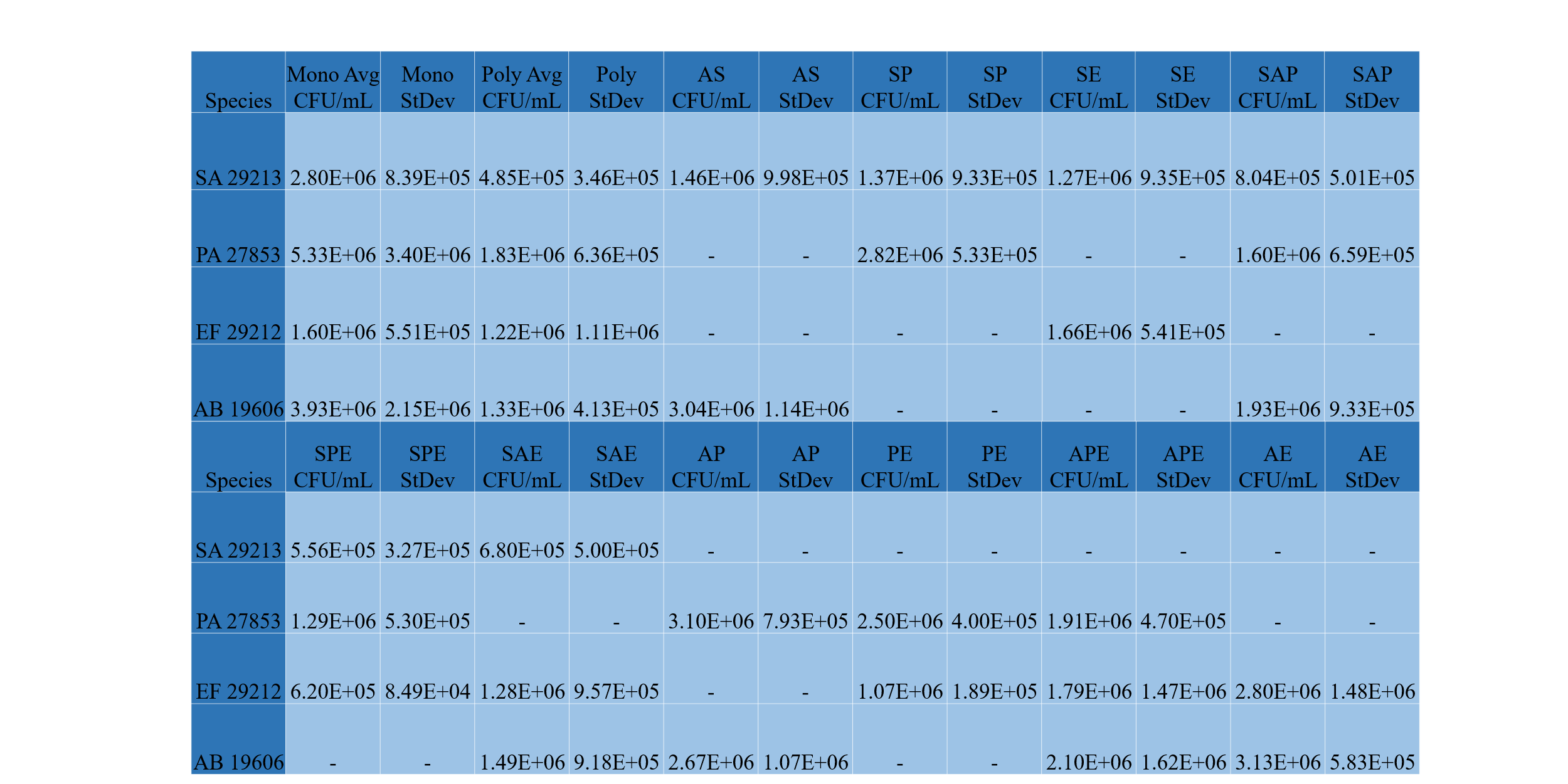

Each organism is abbreviated in the first column by species and genus initial (“SA” for *S.*

*aureus*, “PA” for *P. aeruginosa*, etc.) Each organism within communities is abbreviated in the

first and sixth row by the first initial of each species (“S” for *S. aureus*, “P” for *P. aeruginosa*,

etc.).

**Supplemental Table A2. Inoculum CFU/mL obtained for each species shows inoculums were very similar across species in the polymicrobial condition.^a^**

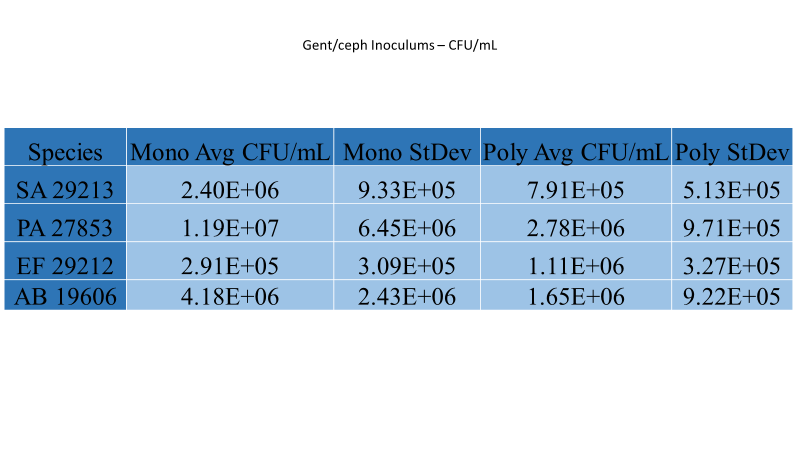

^a^Average CFU/mL for each species’ inoculum for both the monomicrobial and polymicrobial conditions. The standard deviation is also included.

**Supplemental Table A3.** **Observed gentamicin visible turbidity MICs do not necessarily match CFUs obtained.^a^**

| **Organism** | **MIC (Observed Turbidity)** |
| --- | --- |
| *S. aureus* (n=9) | 0.167±0.072 |
| *P. aeruginosa* (n=9) | 0.417±0.144 |
| *E. faecalis* (n=12) | 2.5±1 |
| *A. baumannii* (n=12) | 3.5±1 |
| Polymicrobial (n=9) | 4±0; presumed *P. aeruginosa* MIC 0.5±0 |

^a^All species were within range for their monomicrobial MIC breakpoints for gentamicin. CFU/mL values obtained for each species are shown in Supplemental Figure A1 and display a disparity between where visible breakpoints are read and where CFU counts are no longer detectable. For example, *S. aureus* has measurable levels of growth until a concentration of 2 μg/mL, yet the visible turbidity MIC of gentamicin was read to average at 0.167 μg/mL, a much lower concentration. “Presumed *P. aeruginosa*” MIC is based on the blue-green coloration of pyocyanin present.

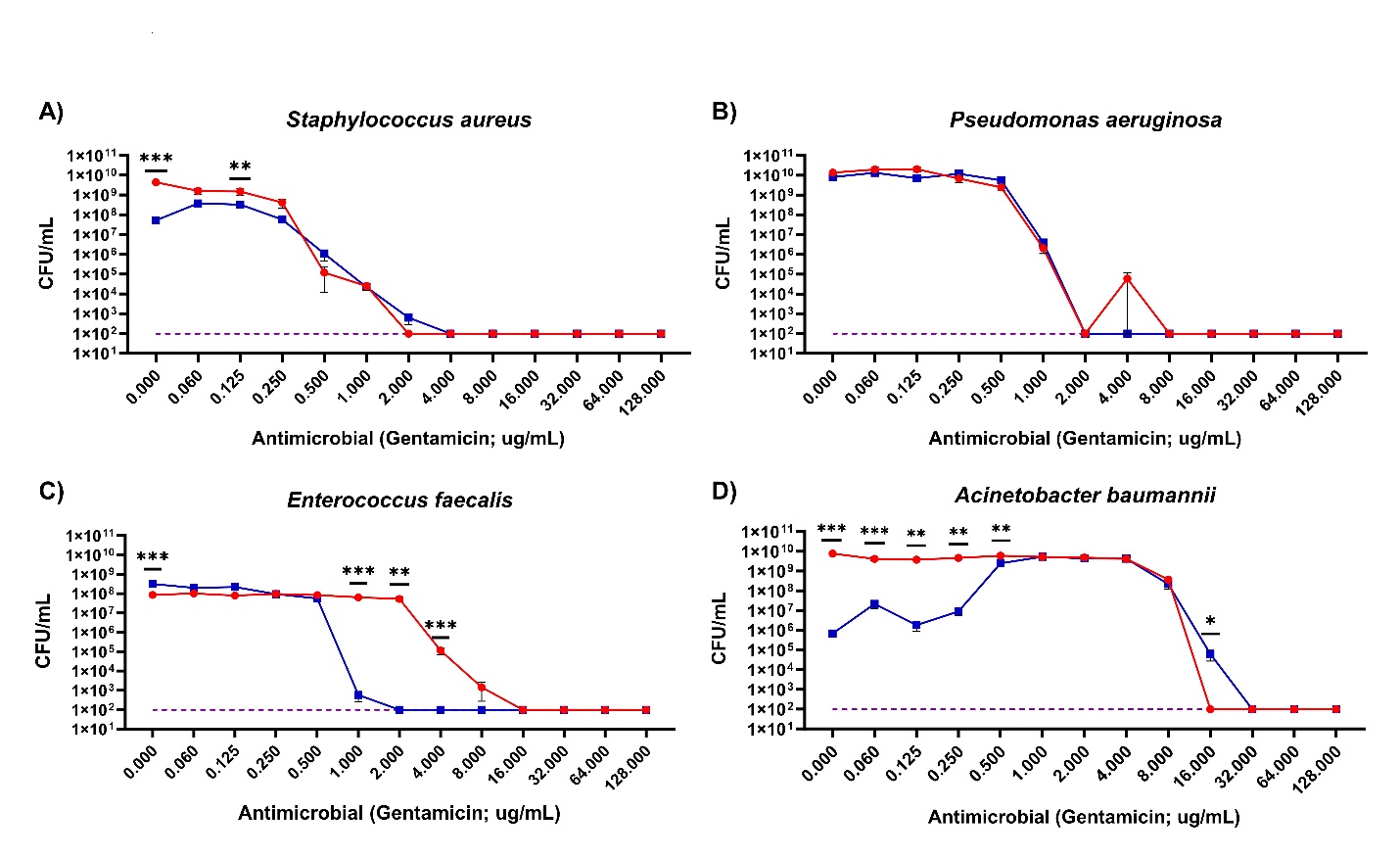

**Supplemental Figure A1. *Enterococcus faecalis* shows increased susceptibility to gentamicin when grown in the polymicrobial condition.** The most striking phenomenon after treatment with gentamicin in the mono versus polymicrobial conditions was the increase in susceptibility to gentamicin of *E. faecalis* from 1 μg/mL to 8 μg/mL. Also worth noting is the decrease in *A. baumannii* growth in the polymicrobial condition due to antagonism by *P. aeruginosa*. As *Pseudomonas* begins to die off, it loses its ability to antagonize its fellow community members, and thus *A. baumannii* begins to make a comeback. All lines are indicative of data averaged from 9-12 technical replicates across 3-4 biological replicates (3 technical replicates per biological) as outlined in Supplemental Table A3. Asterisks represent significance as determined by a Mann Whitney test.

**Supplemental Table A4.** **Observed tetracycline visible turbidity MICs do not necessarily match CFUs obtained.**^a^

| **Organism** | **MIC (Observed Turbidity)** |
| --- | --- |
| *S. aureus* (n=9) | 0.167±0.072 |
| *P. aeruginosa* (n=9) | 0.417±0.144 |
| *E. faecalis* (n=12) | 2.5±1 |
| *A. baumannii* (n=12) | 3.5±1 |
| Polymicrobial (n=9) | 4±0; presumed *P. aeruginosa* MIC 0.5±0 |

^a^All species were within range for their monomicrobial MIC breakpoints for tetracycline. CFU/mL values obtained for each species are shown in Supplemental Figure A2 and display a disparity between where visible breakpoints are read and where CFU counts are no longer detectable. For example, *P. aeruginosa* has measurable levels of growth at all concentrations of tetracycline, yet the visible turbidity MIC of tetracycline was read to average at 13.333 μg/mL, a much lower concentration than the highest concentration tested (128 μg/mL).

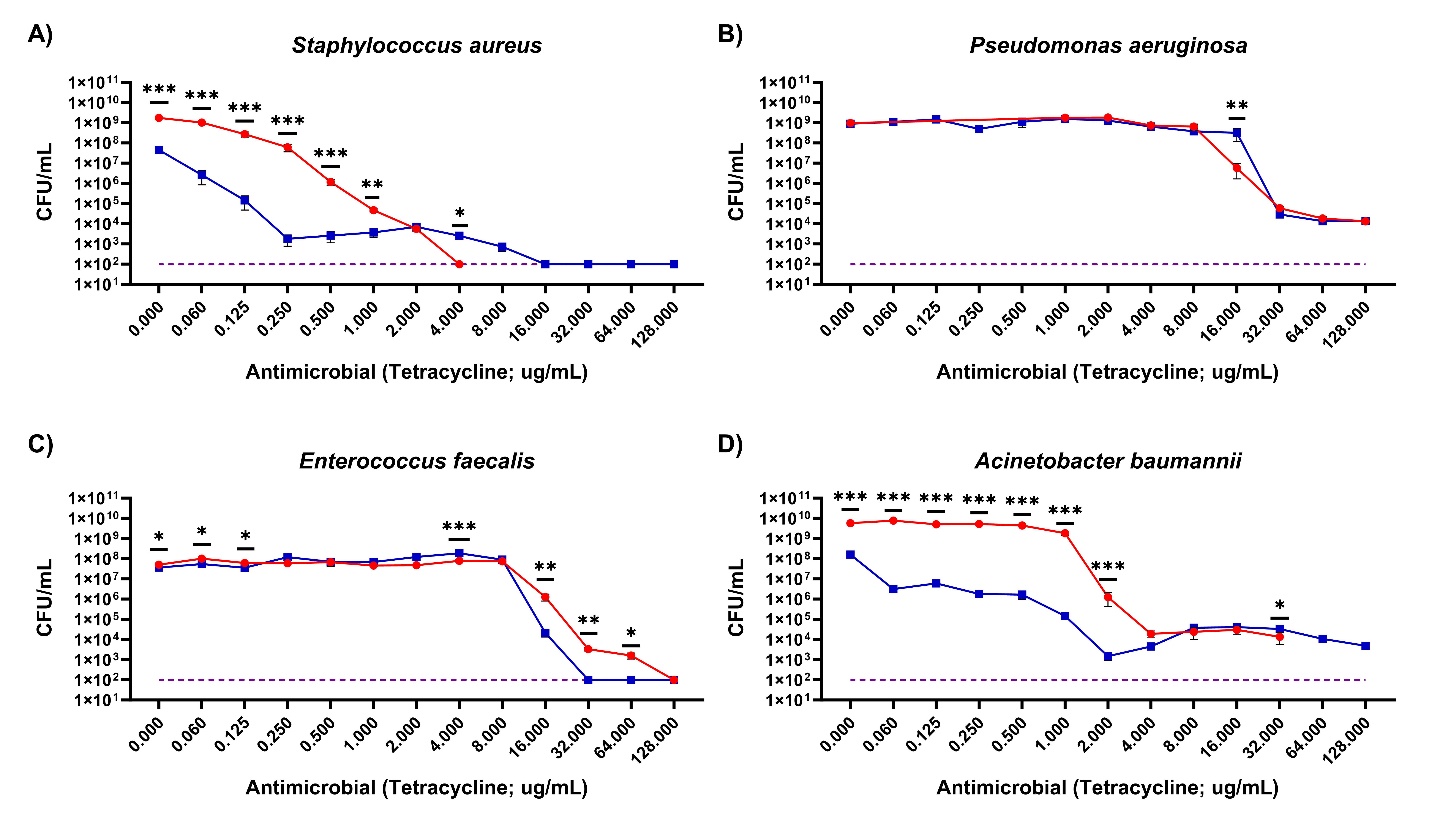

**Supplemental Figure A2.** ***Staphylococcus aureus* shows increased susceptibility to tetracycline at lower concentrations but decreased susceptibility at higher concentrations when grown in the polymicrobial condition.** The most interesting phenomenon after treatment with tetracycline in the mono versus polymicrobial conditions was the increase in susceptibility to tetracycline of *S. aureus* at concentrations ranging from 0.06 μg/mL to 1 μg/mL when grown in the polymicrobial condition. However, at higher concentrations ranging from 4 μg/mL to 16 μg/mL, *S. aureus* showed a decrease in susceptibility, showing better survival of the bacterial in the community. Also worth noting is the decrease in *A. baumannii* growth in the polymicrobial condition at lower concentrations of tetracycline. All lines are indicative of data averaged from 9-12 technical replicates across 3-4 biological replicates (3 technical replicates per biological) as outlined in Supplemental Table A4. Asterisks represent significance as determined by a Mann Whitney test.

**Supplemental Table A5.** **Observed rifampin visible turbidity MICs do not necessarily match CFUs obtained.^a^**

| **Organism** | **MIC (Observed Turbidity)** |
| --- | --- |
| *S. aureus* (n=9) | 0.145±0.097 |
| *P. aeruginosa* (n=9) | 128±0 |
| *E. faecalis* (n=9) | 4±0 |
| *A. baumannii* (n=9) | 16±0 |
| Polymicrobial (n=9) | 106.667±36.95; presumed *P. aeruginosa* MIC 85.333±36.95 |

^a^*E. faecalis* and *A. baumannii* were both within range for their monomicrobial MIC breakpoints for rifampin. *S. aureus* had one run in which its MIC breakpoint was too high, and *P. aeruginosa*’s breakpoint was too high for all three runs. CFU/mL values obtained for each species are shown in Supplemental Figure A3 and display a disparity between where visible breakpoints are read and where CFU counts are no longer detectable. For every species except *S. aureus*, there was a detectable level of monomicrobial growth for all concentrations of rifampin up to 128 μg/mL. Yet even *Pseudomonas*, with its breakpoint recorded at 128 μg/mL, was supposedly inhibited by rifampin, at least when read by visible turbidity. The other species had breakpoints much lower, and yet still had CFUs at all concentrations of rifampin tested.

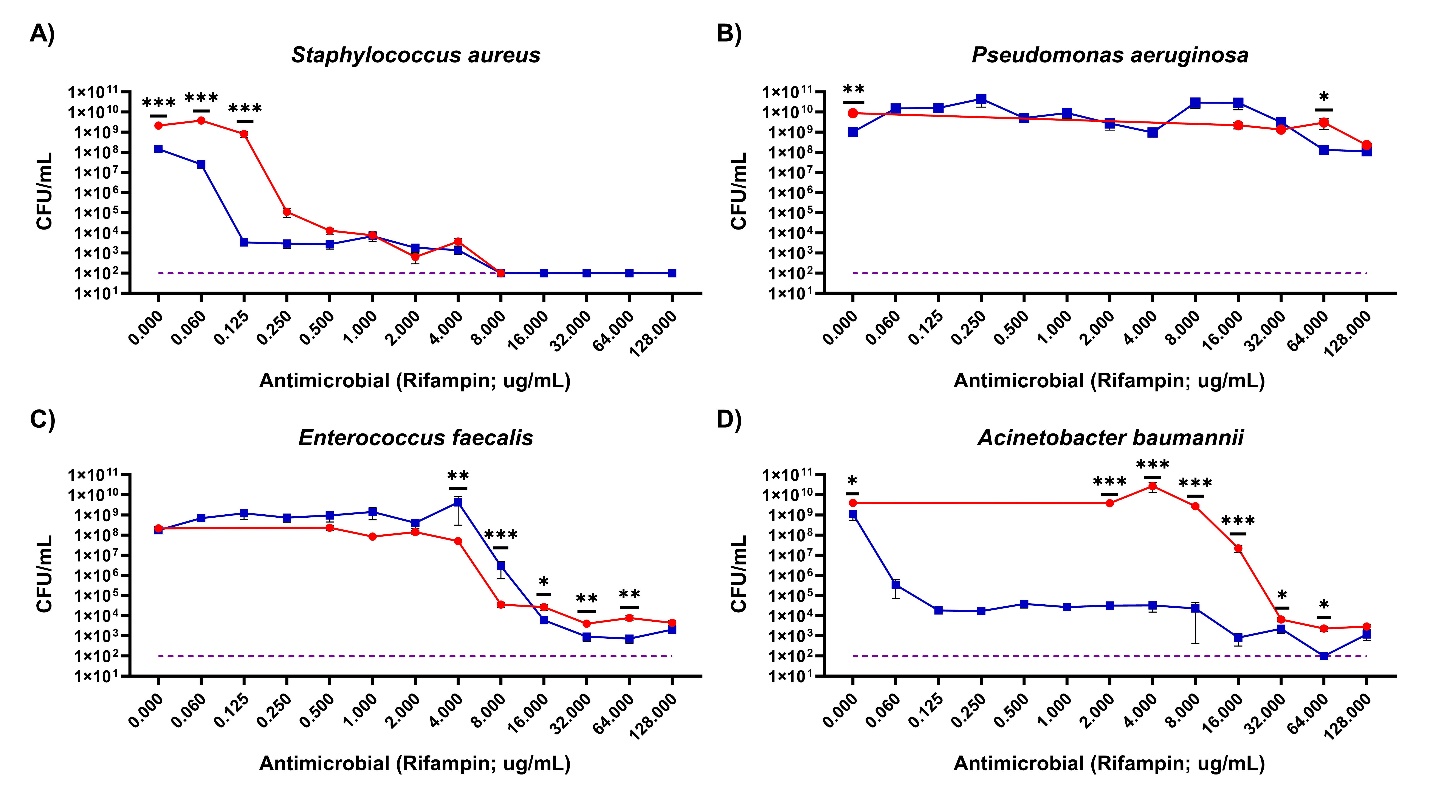

**Supplemental Figure A3. Both** ***Staphylococcus aureus* and *Acinetobacter baumannii* show increased susceptibility to rifampin when grown in the polymicrobial condition.** The most interesting phenomenon after treatment with rifampin in the mono versus polymicrobial conditions was the increase in susceptibility to rifampin of *S. aureus* at concentrations ranging from 0.06 μg/mL to 0.5 μg/mL and of *A. baumannii* at concentrations ranging from 2 μg/mL to 128 μg/mL (2 μg/mL was the cut off for plating, so differences below that were not measured). All lines are indicative of data averaged from 9 technical replicates across 3 biological replicates (3 technical replicates per biological). Asterisks represent significance as determined by a Mann Whitney test.

**Supplemental Table A6.** **Observed ceftazidime visible turbidity MICs do not necessarily match CFUs obtained.**^a^

| **Organism** | **MIC (Observed Turbidity)** |
| --- | --- |
| *S. aureus* (n=9) | 42.667±18.475 |
| *P. aeruginosa* (n=9) | 16±0 |
| *E. faecalis* (n=9) | 8±0 |
| *A. baumannii* (n=9) | 32±0 |
| Polymicrobial (n=9) | 32±0; presumed *P. aeruginosa* MIC 8±0 |

^a^The MIC breakpoints for all four species were too high for all three runs. CFU/mL values obtained for each species are shown in Supplemental Figure A4 and display a disparity between where visible breakpoints are read and where CFU counts are no longer detectable. The most striking difference between visible turbidity and CFU counts is with *E. faecalis*. *E. faecalis* maintains a high CFU/mL concentration throughout all concentrations of ceftazidime, and yet has a visible turbidity MIC breakpoint of 8 μg/mL.

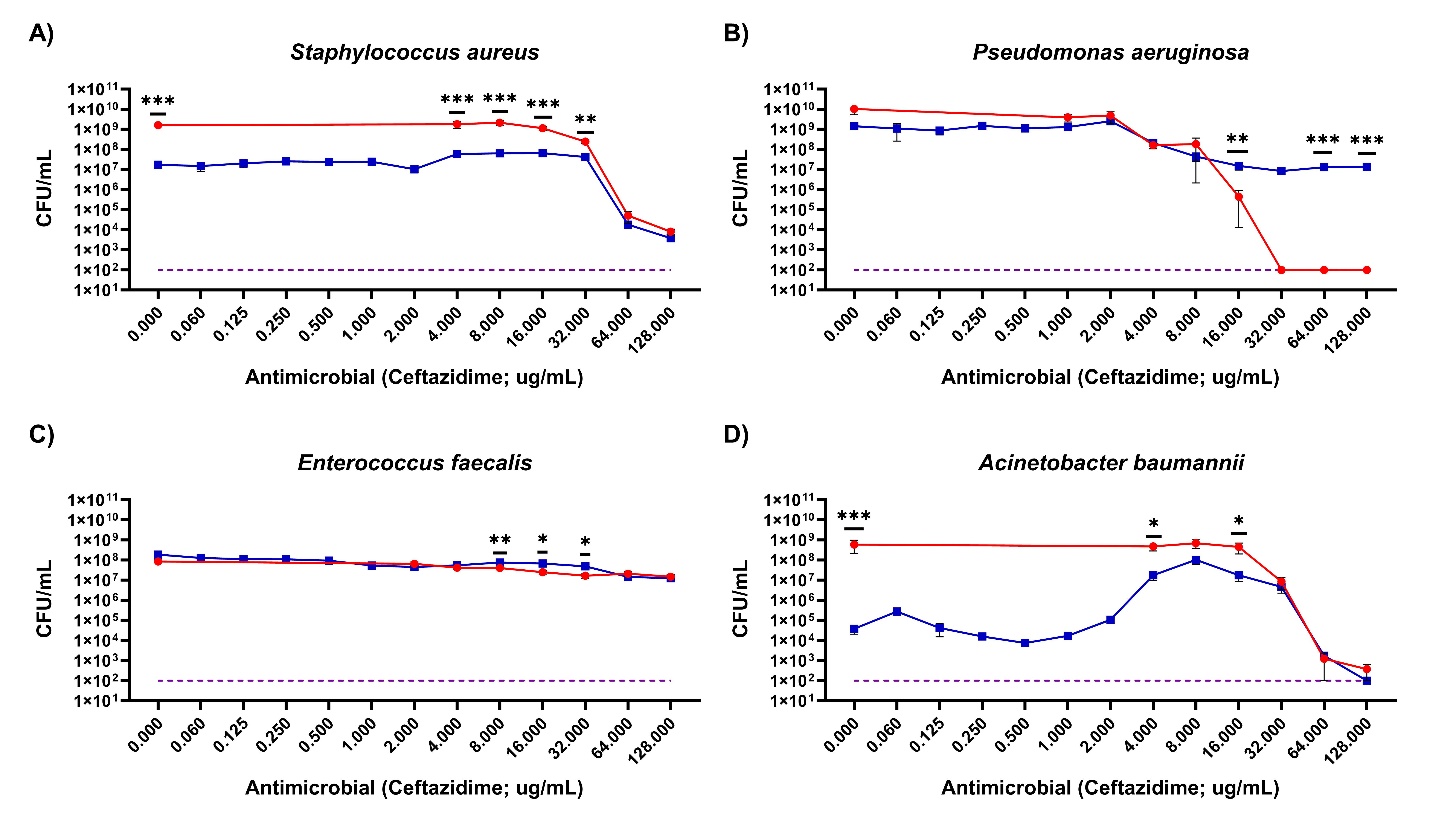

**Supplemental Figure A4. *P. aeruginosa* shows decreased susceptibility to ceftazidime when grown in the polymicrobial condition.** The most interesting phenomenon after treatment with ceftazidime in the mono versus polymicrobial conditions was the decrease in susceptibility to ceftazidime of *P. aeruginosa* at concentrations ranging from 32 μg/mL to 128 μg/mL. This shows increased survival when *Pseudomonas* is in a community. All lines are indicative of data averaged from 9 technical replicates across 3 biological replicates (3 technical replicates per biological). Asterisks represent significance as determined by a Mann Whitney test.

**Supplemental Table A7.** **Observed cephalexin visible turbidity MICs do not necessarily match CFUs obtained.**^a^

| **Organism** | **MIC (Observed Turbidity)** |
| --- | --- |
| *S. aureus* (n=9) | 2±0 |
| *P. aeruginosa* (n=9) | >128±0 |
| *E. faecalis* (n=9) | 53.333±18.475 |
| *A. baumannii* (n=9) | >128±0 |
| Polymicrobial (n=9) | >128±0; presumed *P. aeruginosa* MIC >128±0 |

^a^There are no published breakpoints for any of these 4 species in the CLSI manual. CFU/mL values obtained for each species are shown in Supplemental Figure A5 and display a disparity between where visible breakpoints are read and where CFU counts are no longer detectable. The most striking difference between visible turbidity and CFU counts is with *S. aureus*. *S. aureus* begins to decline in the number of CFU/mL, but then starts increasing again at 8 μg/mL. Yet the MIC breakpoint for *S. aureus* in the monomicrobial condition is read as 2 μg/mL.

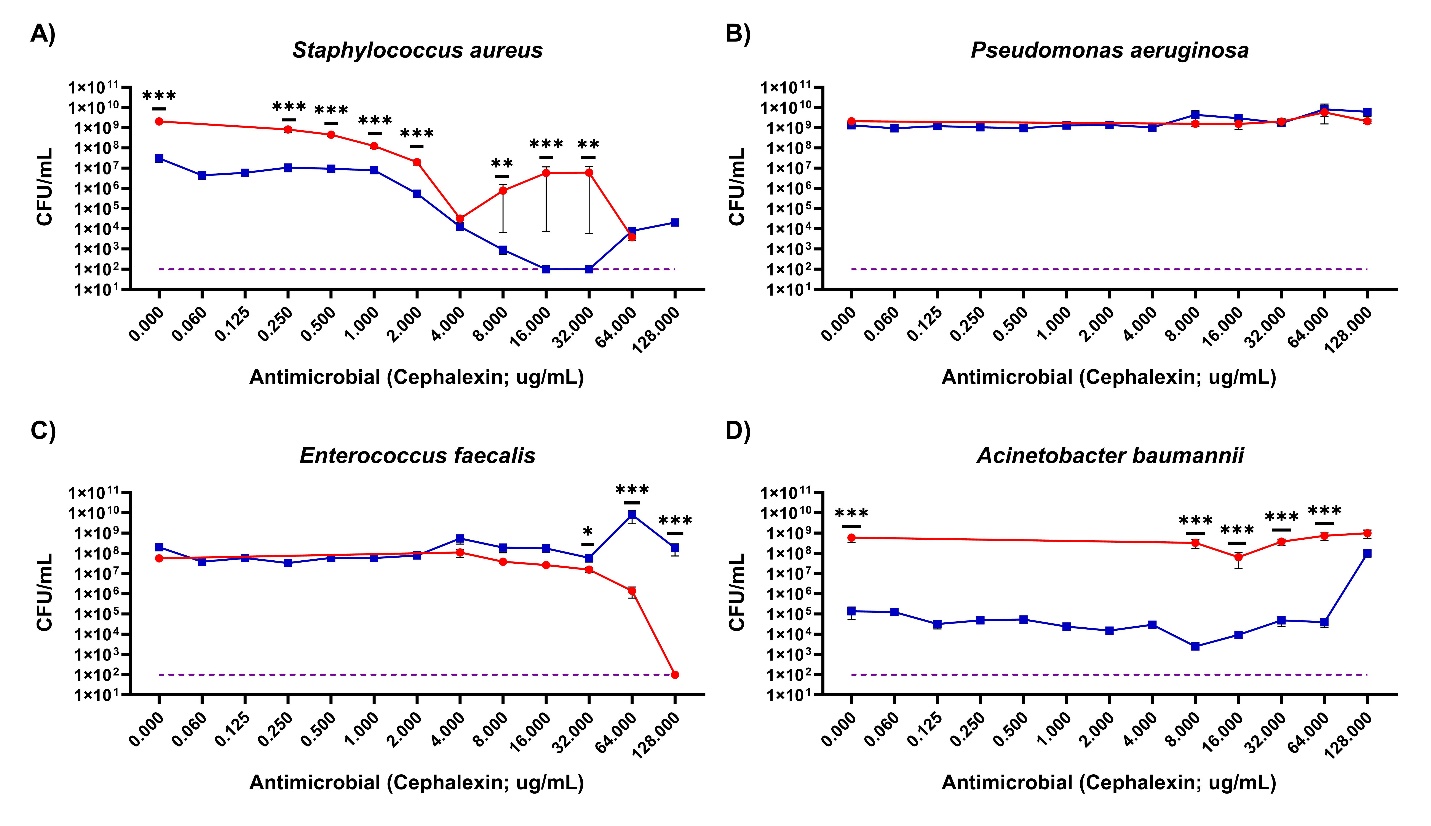

**Supplemental Figure A5. *Enterococcus faecalis* shows decreased susceptibility to cephalexin when grown in the polymicrobial condition, while both *Staphylococcus aureus* and *Acinetobacter baumannii* show increased susceptibility.** The most interesting phenomenon after treatment with cephalexin in the mono versus polymicrobial conditions was the decrease in susceptibility to cephalexin of *E. faecalis* at concentrations ranging from 4 μg/mL to 128 μg/mL (the monomicrobial data for antibiotic concentrations less than 4 μg/mL were not plated to save plates, and thus there is no measured comparable data for these points). Both *S. aureus* and *A. baumannii* had increased susceptibility to cephalexin at all comparable concentrations, showing that while the community helps *E. faecalis* to survive, antagonism hurts both *S. aureus* and *A,* baumannii. All lines are indicative of data averaged from 9 technical replicates across 3 biological replicates (3 technical replicates per biological). Asterisks represent significance as determined by a Mann Whitney test.

**Supplemental Table A8.** **Observed ciprofloxacin visible turbidity MICs do not necessarily match CFUs obtained.^a^**

| **Organism** | **MIC (Observed Turbidity)** |
| --- | --- |
| *S. aureus* (n=9) | 0.25±0 |
| *P. aeruginosa* (n=9) | 0.125±0 |
| *E. faecalis* (n=9) | 0.5±0 |
| *A. baumannii* (n=9) | 0.5±0 |
| Polymicrobial (n=9) | 0.5±0; presumed *P. aeruginosa* MIC <0.06±0 |

^a^All four species had MIC breakpoints within the published ranges in the CLSI manual. CFU/mL values obtained for each species are shown in Supplemental Figure A6 and display a disparity between where visible breakpoints are read and where CFU counts are no longer detectable. The most striking difference between visible turbidity and CFU counts is with *S. aureus*. *S. aureus* never drops below detectable CFU counts up to the last monomicrobial concentration of antibiotic I plated, which was 8 μg/mL. Yet the MIC breakpoint for *S. aureus* in the monomicrobial condition is read as 0.25 μg/mL.

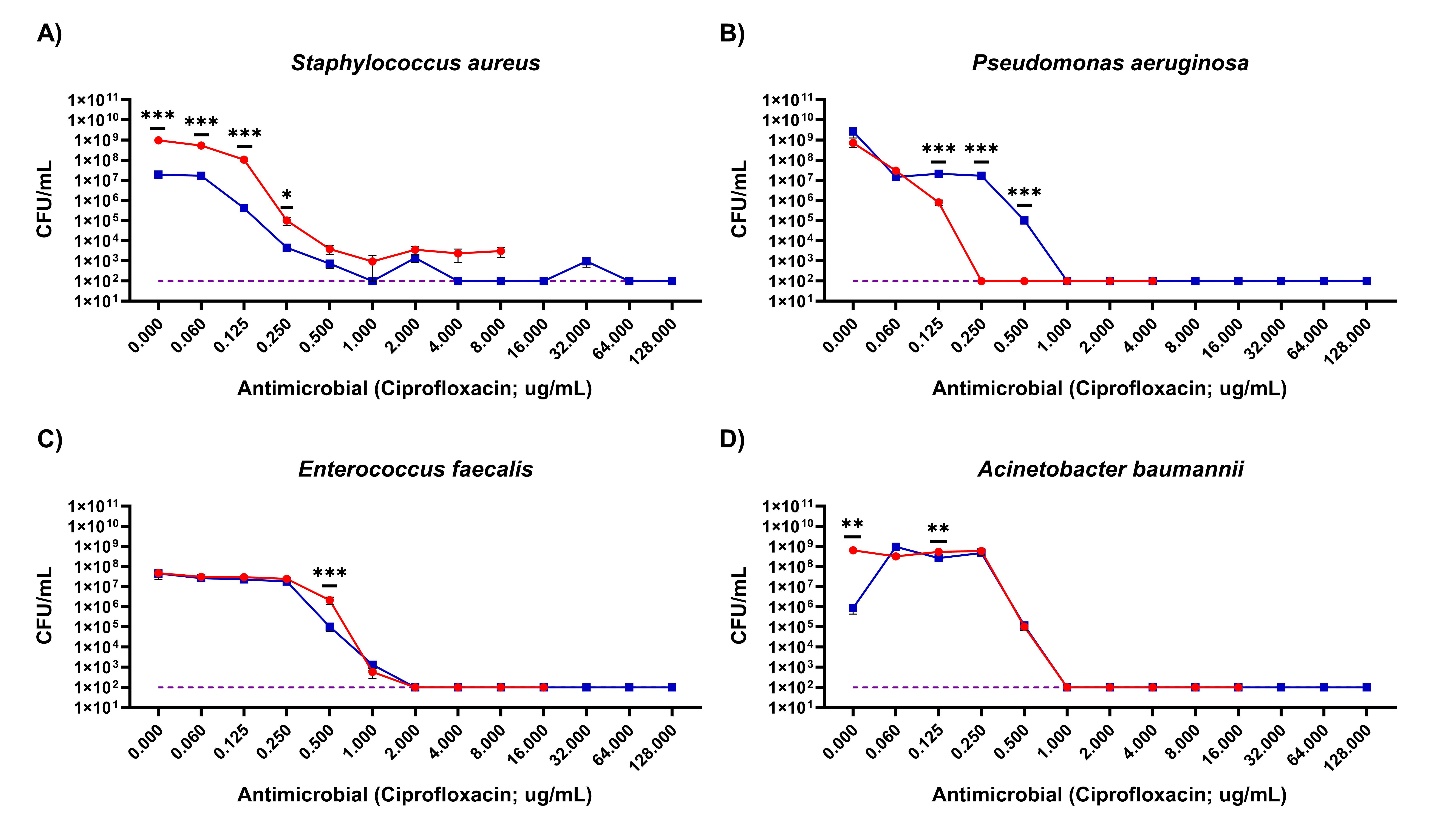

**Supplemental Figure A6. *Pseudomonas aeruginosa* shows decreased susceptibility to ciprofloxacin when grown in the polymicrobial condition, while *Staphylococcus aureus* shows increased susceptibility.** The most interesting phenomenon after treatment with ciprofloxacin in the mono versus polymicrobial conditions was the decrease in susceptibility to ciprofloxacin of *P. aeruginosa* at concentrations ranging from 0.125 μg/mL to 1 μg/mL. *S. aureus* had increased susceptibility to ciprofloxacin at all comparable concentrations. All lines are indicative of data averaged from 9 technical replicates across 3 biological replicates (3 technical replicates per biological). Asterisks represent significance as determined by a Mann Whitney test.

**Supplemental Table A9.** **Observed augmentin visible turbidity MICs do not necessarily match CFUs obtained.^a^**

| **Organism** | **MIC (Observed Turbidity)** |
| --- | --- |
| *S. aureus* (n=9) | 1.167±0.764 |
| *P. aeruginosa* (n=9) | >128±0 |
| *E. faecalis* (n=9) | 0.667±0.289 |
| *A. baumannii* (n=9) | >128±110.851 |
| Polymicrobial (n=9) | >128±0; presumed *P. aeruginosa* MIC >128±0 |

^a^*S. aureus*’ breakpoint was higher than reported in the CLSI manual, while *E. faecalis*’ was in range. Neither *P. aeruginosa* nor *A. baumannii* have breakpoints published in the CLSI manual for augmentin. CFU/mL values obtained for each species are shown in Supplemental Figure A7 and display a disparity between where visible breakpoints are read and where CFU counts are no longer detectable. Notably, *S. aureus* has detectable CFU counts at concentrations up to 64 μg/mL, yet the MIC determined by visible turbidity is 1.167 μg/mL.

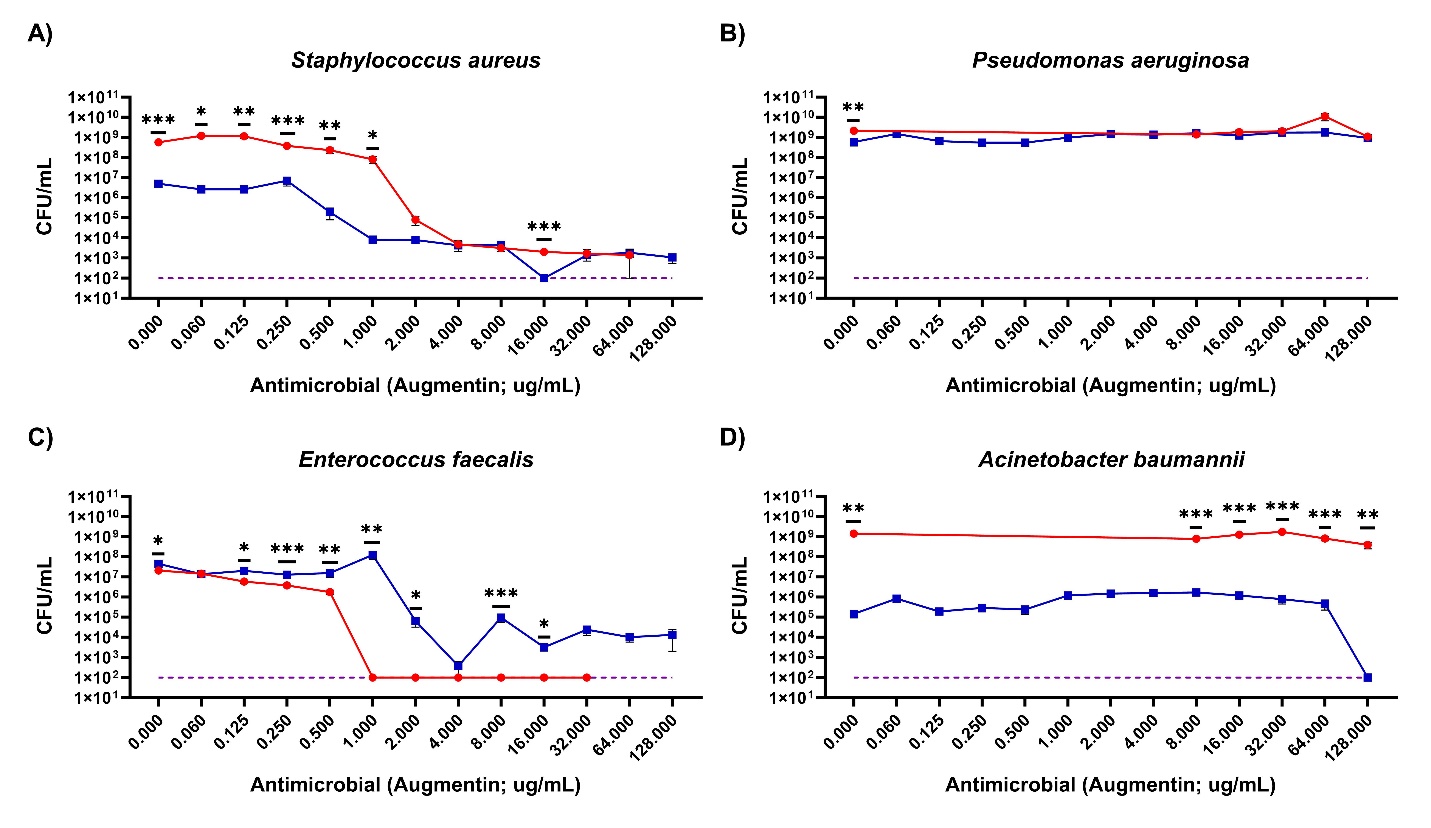

**Supplemental Figure A7. *Enterococcus faecalis* shows decreased susceptibility to augmentin when grown in the polymicrobial condition, while *Staphylococcus aureus* shows increased susceptibility.** The most striking phenomenon after treatment with augmentin in the mono versus polymicrobial conditions was the decrease in susceptibility to augmentin of *E. faecalis* at all measured concentrations. *S. aureus* had increased susceptibility to augmentin at almost all comparable concentrations. All lines are indicative of data averaged from 9 technical replicates across 3 biological replicates (3 technical replicates per biological). Asterisks represent significance as determined by a Mann Whitney test.

**Supplemental Table A10.** **Observed vancomycin visible turbidity MICs do not necessarily match CFUs obtained.^a^**

| **Organism** | **MIC (Observed Turbidity)** |
| --- | --- |
| *S. aureus* (n=9) | 0.194±0.120 |
| *P. aeruginosa* (n=9) | >128±0 |
| *E. faecalis* (n=9) | 0.667±0.289 |
| *A. baumannii* (n=9) | 32±0 |
| Polymicrobial (n=9) | >128±0; presumed *P. aeruginosa* MIC >128±0 |

^a^Both *S. aureus* and *E. faecalis*’ breakpoints were lower than reported in the CLSI manual. Neither *P. aeruginosa* nor *A. baumannii* have breakpoints published in the CLSI manual for vancomycin. CFU/mL values obtained for each species are shown in Supplemental Figure A8 and display a disparity between where visible breakpoints are read and where CFU counts are no longer detectable. Notably, *S. aureus* has detectable CFU counts at concentrations up to 1 μg/mL, yet the MIC determined by visible turbidity is 0.194 μg/mL.

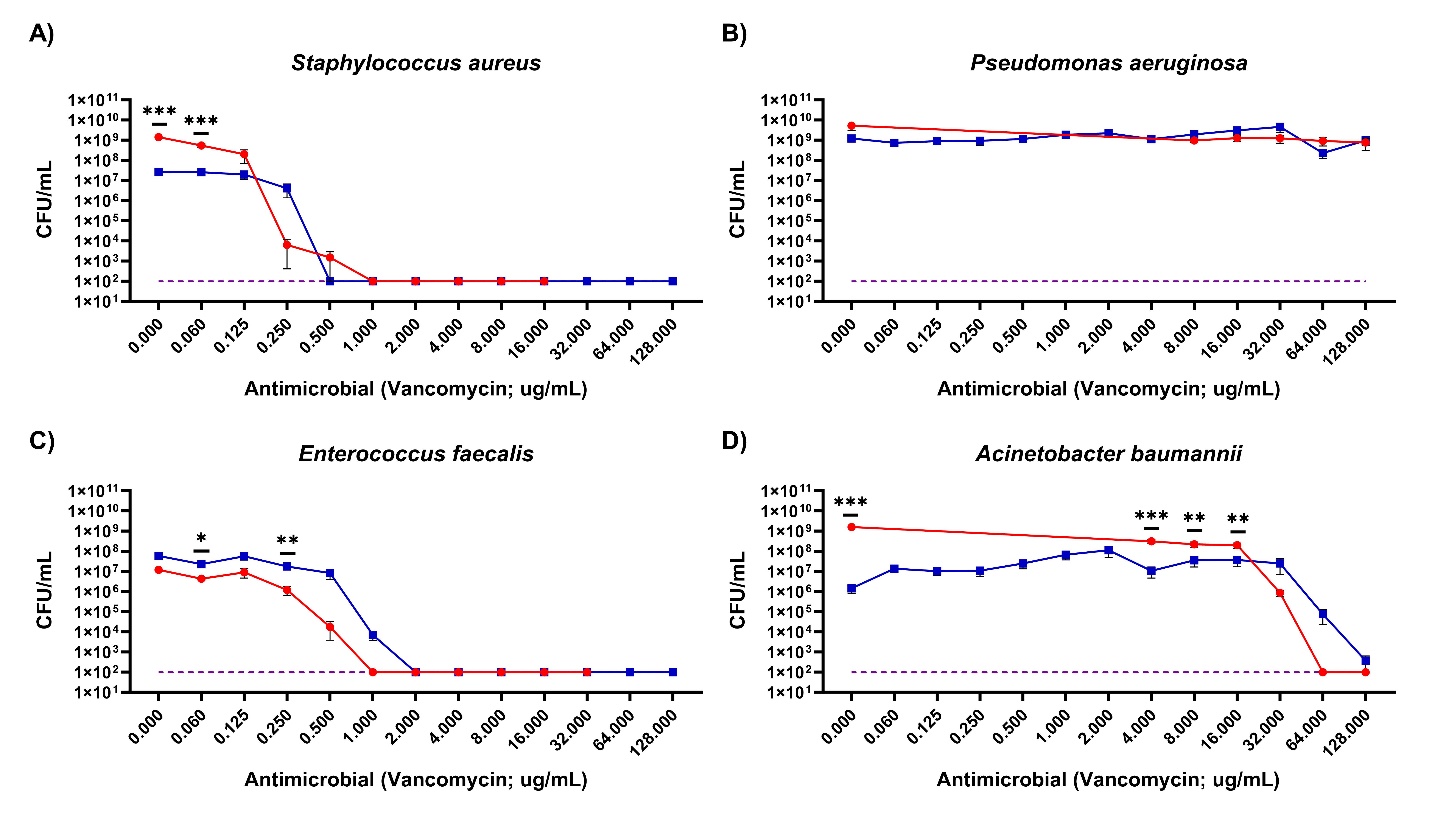

**Supplemental Figure A8. *Enterococcus faecalis* shows decreased susceptibility to vancomycin when grown in the polymicrobial condition.** The most striking phenomenon after treatment with vancomycin in the mono versus polymicrobial conditions was the decrease in susceptibility to vancomycin of *E. faecalis* at all measured concentrations. All lines are indicative of data averaged from 9 technical replicates across 3 biological replicates (3 technical replicates per biological). Asterisks represent significance as determined by a Mann Whitney test.

**Supplemental Table A11.** **Observed polymyxin B visible turbidity MICs do not necessarily match CFUs obtained.^a^**

| **Organism** | **MIC (Observed Turbidity)** |
| --- | --- |
| *S. aureus* (n=9) | 64±0 |
| *P. aeruginosa* (n=9) | 49.333±68.391 |
| *E. faecalis* (n=9) | 128±0 |
| *A. baumannii* (n=9) | 20.89±14.868 |
| Polymicrobial (n=9) | 85.333±36.95; presumed *P. aeruginosa* MIC 4.443±0.768 |

^a^Both *P. aeruginosa*’s and *A. baumanii*’s breakpoints were higher than reported in the CLSI manual. Neither *S. aureus* nor *E. faecalis* have breakpoints published in the CLSI manual for vancomycin. CFU/mL values obtained for each species are shown in Supplemental Figure A9 and display a disparity between where visible breakpoints are read and where CFU counts are no longer detectable. Notably, *P. aeruginosa* has detectable CFU counts at concentrations up to 64 μg/mL, yet the MIC determined by visible turbidity is 49.333 μg/mL.

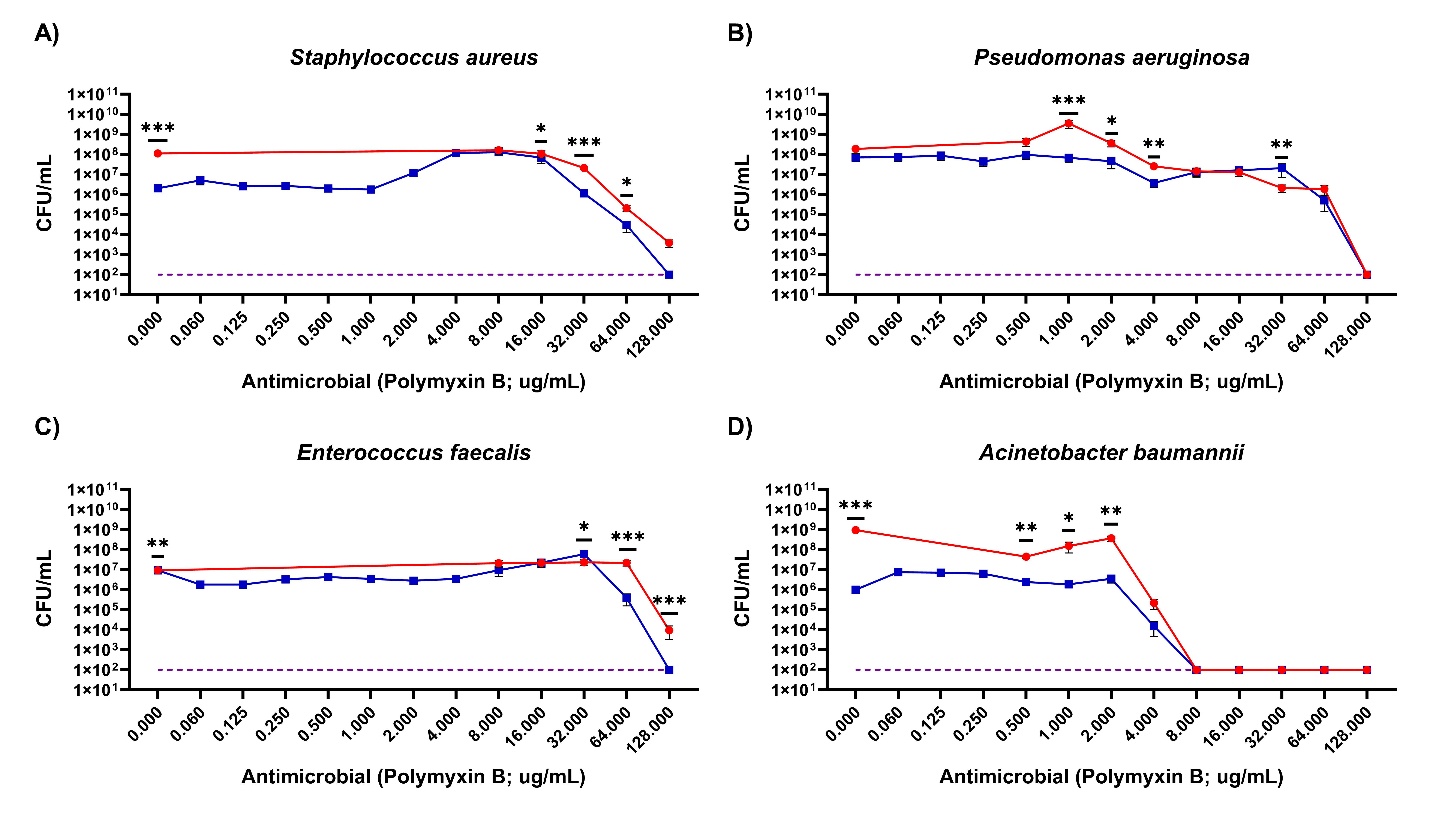

**Supplemental Figure A9. *Enterococcus faecalis* and *Staphylococcus aureus* show increased susceptibility to polymyxin B when grown in the polymicrobial condition.** The most striking phenomenon after treatment with polymyxin B in the mono versus polymicrobial conditions was the increase in susceptibility to polymyxin B of *E. faecalis* and *S. aureus* at 128 µg/mL. All lines are indicative of data averaged from 9 technical replicates across 3 biological replicates (3 technical replicates per biological). Asterisks represent significance as determined by a Mann Whitney test.

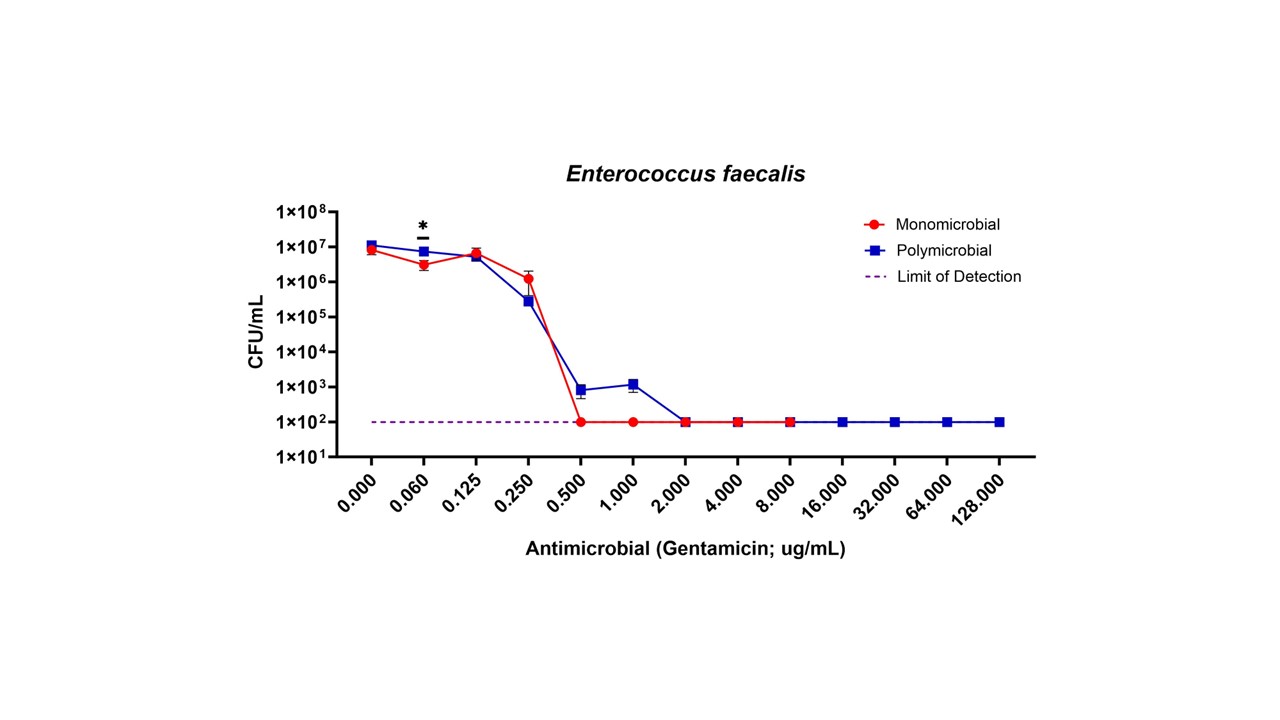

**Supplemental Figure A10. *Enterococcus faecalis*’ increased susceptibility to gentamicin is only present in aerobic conditions and not anaerobic.** *E. faecalis* when grown in an anaerobic environment no longer displayed increased susceptibility to gentamicin when grown in the polymicrobial community. All lines are indicative of data averaged from 12 technical replicates across 4 biological replicates (3 technical replicates per biological). Asterisks represent significance as determined by a Mann Whitney test.

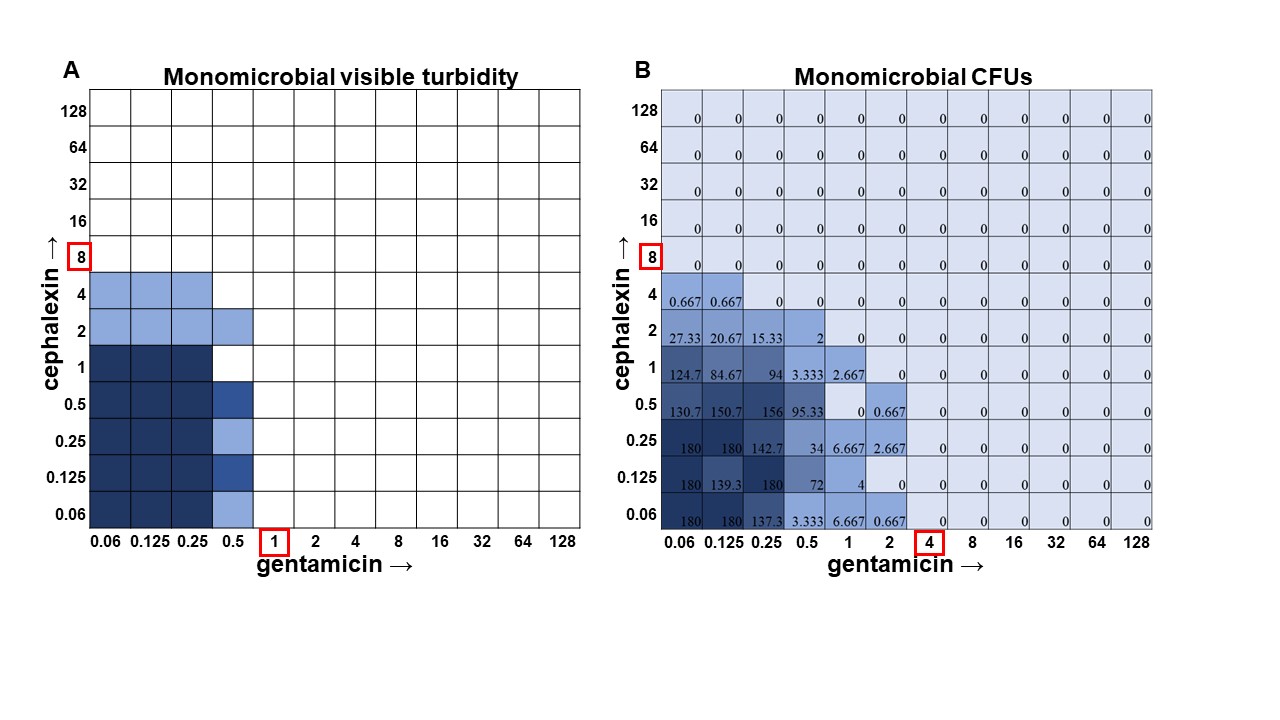

**Supplemental Figure A11.** **Turbidity versus CFU data in *Staphylococcus aureus* checkerboard assay.**

**A.** Visible turbidity of *S. aureus* ATCC 29213 when grown in the monomicrobial condition. Data represents triplicate checkerboard assays with cephalexin and gentamicin. Dark coloration indicates that turbidity was visible in the well in all replicates, medium coloration indicates that turbidity was visible in 2 replicates, light coloration indicates that turbidity was visible in 1 replicate. **B.** CFU data from the same triplicate plates with numbers representing the average of the triplicate CFUs detected. Wells displaying TNTC (too numerous to count) CFUs were assigned a number of 180 to enable the determination of averages across replicates.

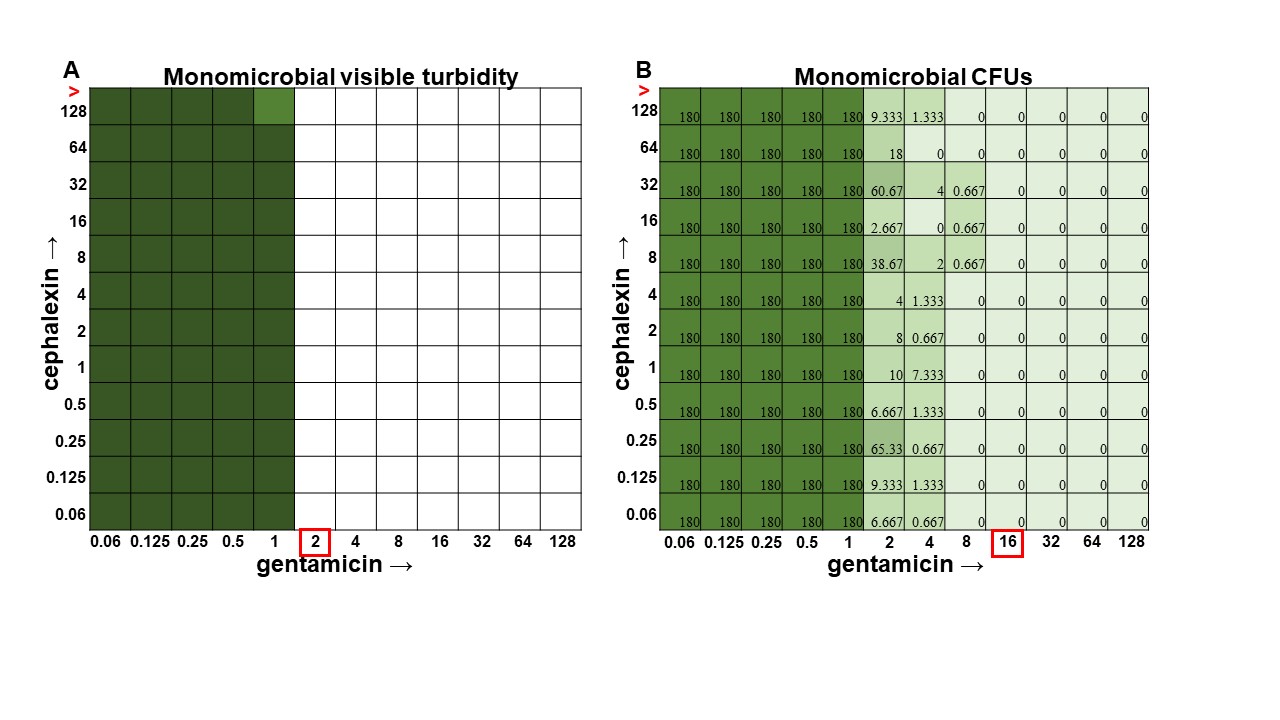

**Supplemental Figure A12.** **Turbidity versus CFU data in *Pseudomonas aeruginosa* checkerboard assay.**

**A.** Visible turbidity of *Pseudomonas aeruginosa* ATCC 27853 when grown in the monomicrobial condition. Data represents triplicate checkerboard assays with cephalexin and gentamicin. Dark coloration indicates that turbidity was visible in the well in all replicates, medium coloration indicates that turbidity was visible in 2 replicates, light coloration indicates that turbidity was visible in 1 replicate. **B.** CFU data from the same triplicate plates with numbers representing the average of the triplicate CFUs detected. Wells displaying TNTC (too numerous to count) CFUs were assigned a number of 180 to enable the determination of averages across replicates.

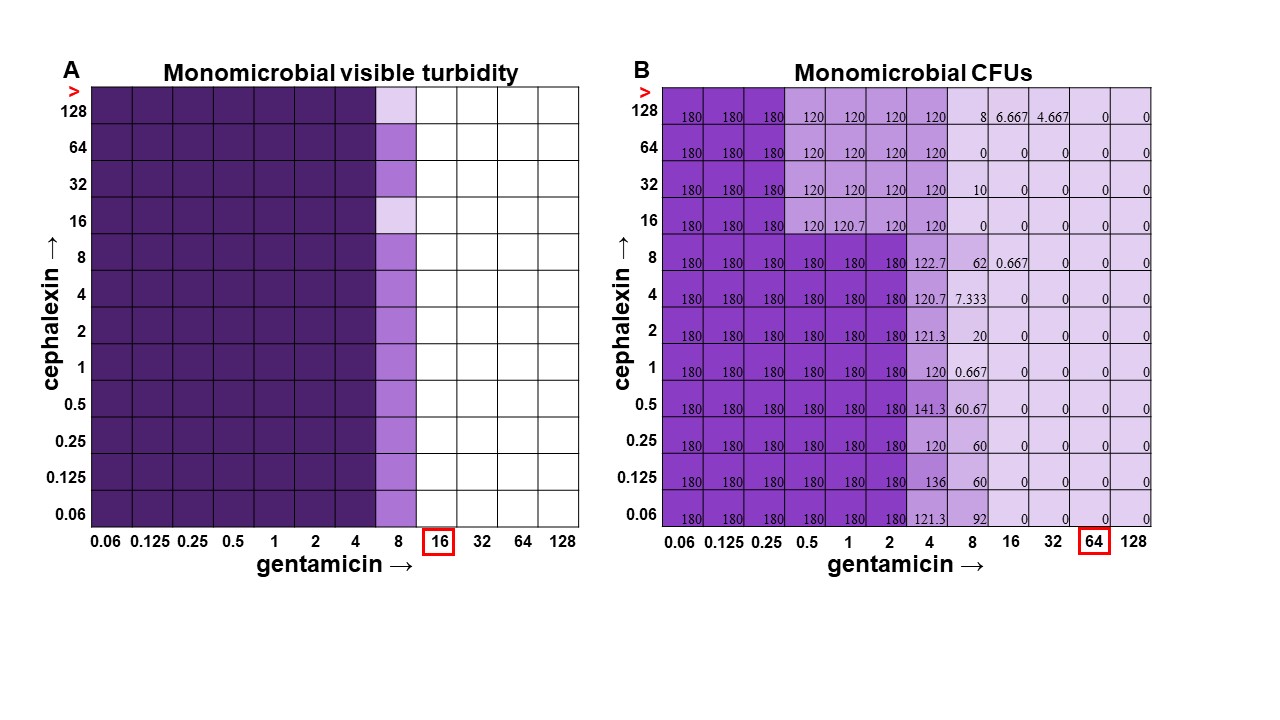

**Supplemental Figure A13.** **Turbidity versus CFU data in *Acinetobacter baumannii* checkerboard assay.**

**A.** Visible turbidity of *Acinetobacter baumannii* ATCC 19606 when grown in the monomicrobial condition. Data represents triplicate checkerboard assays with cephalexin and gentamicin. Dark coloration indicates that turbidity was visible in the well in all replicates, medium coloration indicates that turbidity was visible in 2 replicates, light coloration indicates that turbidity was visible in 1 replicate. **B.** CFU data from the same triplicate plates with numbers representing the average of the triplicate CFUs detected. Wells displaying TNTC (too numerous to count) CFUs were assigned a number of 180 to enable the determination of averages across replicates.

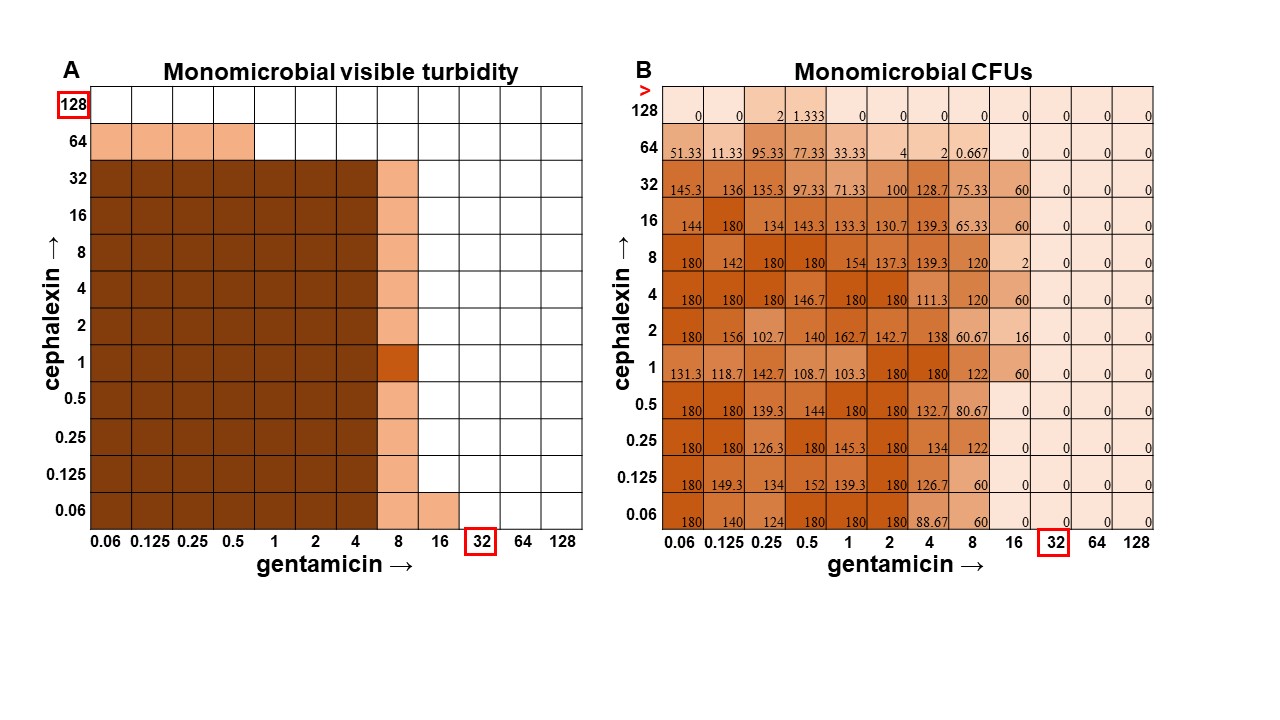

**Supplemental Figure A14. Turbidity versus CFU data in *Enterococcus faecalis* checkerboard assay.**

**A.** Visible turbidity of *Enterococcus faecalis* ATCC 29212 when grown in the monomicrobial condition. Data represents triplicate checkerboard assays with cephalexin and gentamicin. Dark coloration indicates that turbidity was visible in the well in all replicates, medium coloration indicates that turbidity was visible in 2 replicates, light coloration indicates that turbidity was visible in 1 replicate. **B.** CFU data from the same triplicate plates with numbers representing the average of the triplicate CFUs detected. Wells displaying TNTC (too numerous to count) CFUs were assigned a number of 180 to enable the determination of averages across replicates.

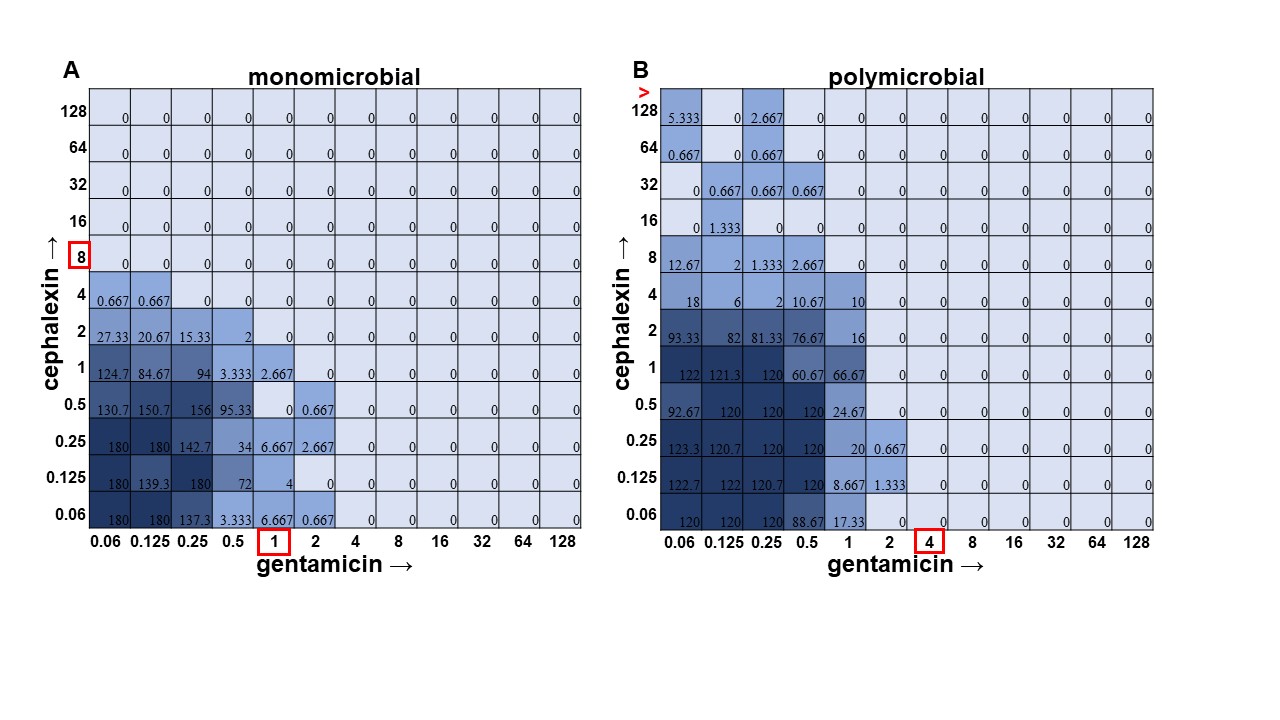

**Supplemental Figure A15.** **Monomicrobial versus polymicrobial checkerboards reveal that *S. aureus* loses sensitivity to cephalexin in polymicrobial culture.** CFU counts for *Staphylococcus aureus* ATCC 29213 when grown in the **A.** monomicrobial condition versus the **B.** polymicrobial condition. As shown above, *S. aureus* becomes less susceptible to cephalexin, growing even at concentrations of 128 μg/mL. Stars represent significance as determined by an unpaired t-test with Welch correction.

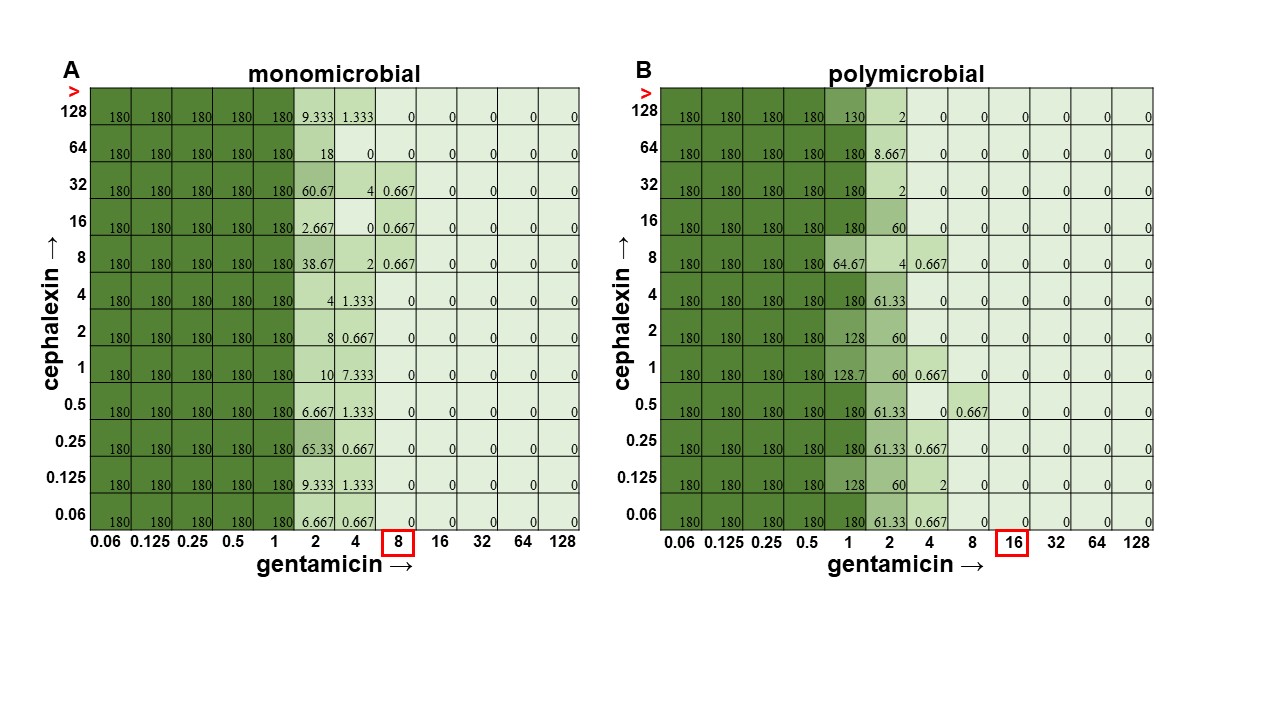

**Supplemental Figure A16. Monomicrobial versus polymicrobial checkerboards show little changes in susceptibility to either gentamicin or cephalexin with *Pseudomonas aeruginosa*.** CFU counts for *P. aeruginosa* ATCC 27853 when grown in the **A.** monomicrobial condition versus the **B.** polymicrobial condition. As shown above, *P. aeruginosa* displays very few changes in susceptibility to either antibiotic. Stars represent significance as determined by an unpaired t-test with Welch correction.

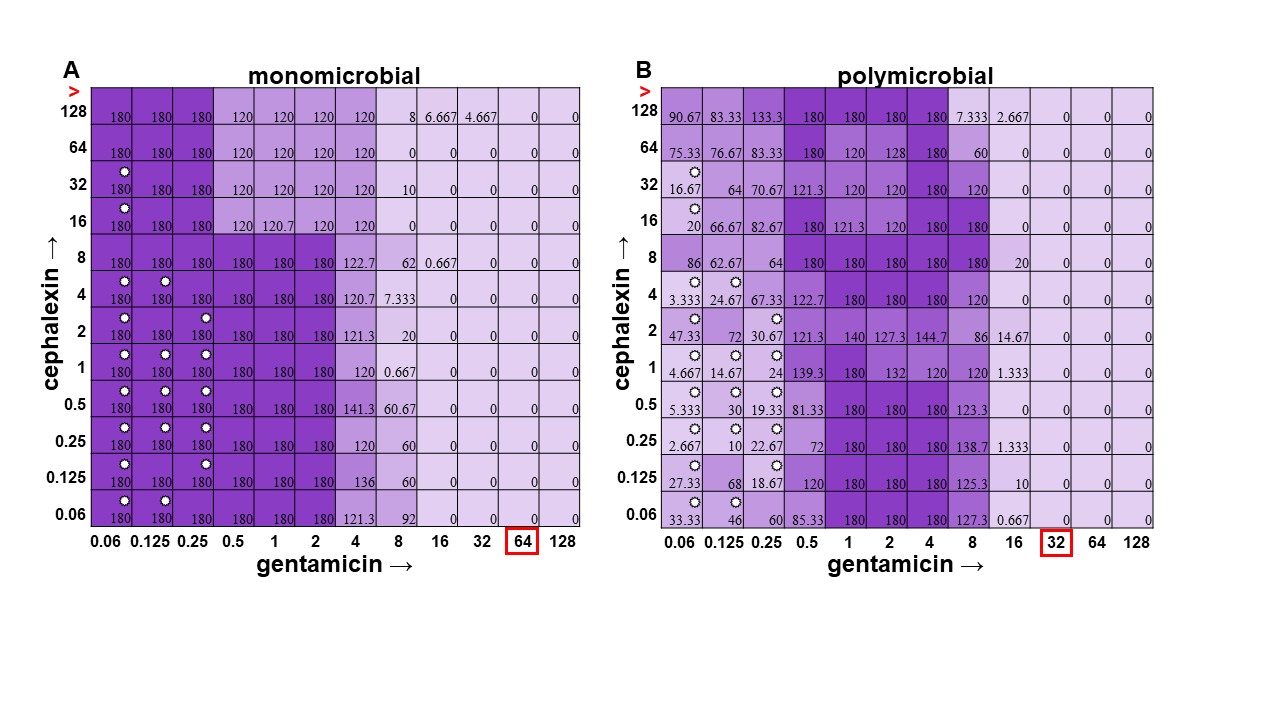

**Supplemental Figure A17. Monomicrobial versus polymicrobial checkerboards reveal that *A. baumannii* succumbs to interspecies competition in low levels of antibiotic but overall MICs are unchanged.** CFU counts for *A. baumannii* ATCC 19606 when grown in the **A.** monomicrobial condition versus the **B.** polymicrobial condition. As shown above, *A. baumannii* growth is inhibited in the polymicrobial condition where *P. aeruginosa* growth is present. This is due to antagonization by its competitor. Stars represent significance as determined by an unpaired t-test with Welch correction.
